# Evolutionary instability of collateral susceptibility networks in ciprofloxacin resistant clinical *Escherichia coli* strains

**DOI:** 10.1101/2021.10.26.465989

**Authors:** Vidar Sørum, Emma L. Øynes, Anna S. Møller, Klaus Harms, Ørjan Samuelsen, Nicole L. Podnecky, Pål J. Johnsen

## Abstract

Collateral sensitivity and resistance occur when resistance development towards one antimicrobial either potentiates or deteriorates the effect of others, respectively. Previous reports on collateral effects on susceptibility focus on newly acquired resistance determinants and propose that novel treatment guidelines informed by collateral networks may reduce the evolution, selection and spread of antimicrobial resistance. In this study, we investigate the evolutionary stability of collateral networks in five ciprofloxacin resistant, clinical *Escherichia coli* strains. After 300 generations of experimental evolution without antimicrobials, we show complete fitness restoration in four of five genetic backgrounds and demonstrate evolutionary instability in collateral networks of newly acquired resistance determinants. We show that compensatory mutations reducing efflux expression is the main driver destabilizing initial collateral networks and identify *rpoS* as a putative target for compensatory evolution. Our results add another layer of complexity to future predictions and clinical application of collateral networks.

## Introduction

The current discovery void in the development of novel antimicrobial agents is at the core of the global antimicrobial resistance crisis. We are at risk of running out of antimicrobials and, consequently, there is an urgent need to prolong the lifespan of existing, effective antimicrobial agents. One promising strategy is to exploit the concepts of collateral sensitivity, and its inverse collateral resistance, where resistance development towards specific antimicrobials modulates susceptibilities towards alternative agents (1). Collateral networks have been proposed for use in antimicrobial cycling protocols (2), sequential treatment regimens (3)(4)(5), and in combination therapies (6)(7) to limit, prevent or reverse antimicrobial resistance evolution, selection and spread (1)(8).

It is clear that future application of collateral effects in antimicrobial resistance management depends on effective and accurate predictions informed by susceptibility testing. Early, seminal work has almost exclusively used emblematic laboratory strains and demonstrated the pervasiveness of collateral networks (2)(9)(10), elucidated mechanistic insights (9)(11), and/or proposed conceptual treatment strategies implementing collateral sensitivity (2)(12)(13). More recently collateral sensitivity was shown to directly affect evolutionary trajectories of resistance in patients during treatment (14) and that principal contributors, including mechanisms of resistance, allow robust predictions of collateral susceptibility networks (4)(15)(16). Taken together, these reports allow for careful optimism for future use of collateral networks as an antimicrobial resistance management strategy. However, with few exceptions (3)(17), the current body of literature rests on newly acquired resistance determinants upon which collateral networks are identified and characterized relative to a susceptible wild-type (WT) in an optimal environment as baseline. Thus, robust predictions of collateral networks are contingent on the evolutionary stability of the initial association between resistance determinants and bacterial hosts. A recent study demonstrated that *Pseudomonas aeruginosa* could not overcome collateral sensitivity by *de novo* mutations when challenged with antimicrobials, suggesting robust collateral networks for specific combinations of antimicrobials (3). However, the effect of co-evolution between the bacterial host and its newly acquired antibiotic resistance determinants on the sign and magnitude of collateral networks is currently unknown.

Here we asked if collateral networks were stable over 300 generations of experimental evolution in the absence of antimicrobial selection. We focus here on five different clinical *Escherichia coli* strains harboring newly acquired ciprofloxacin resistance due to combinations of drug target and efflux mutations. Evolved strains were characterized with respect to their collateral networks, relative fitness measurements, whole genome sequencing, and gene expression analyses. We demonstrate evolutionary instability of collateral networks after experimental evolution independent of resistance mechanisms and fitness compensatory mutations. These results introduce an additional layer of complexity with respect to accurate predictions of collateral networks in clinical application.

## Results

### Collateral networks display evolutionary instability

We previously reported conserved collateral effects in a collection of genetically diverse clinical strains of *E. coli* isolated from urinary tract infections (4). From that study, we chose five laboratory-selected ciprofloxacin resistant (CIP-R) *E. coli* strains (4) covering a diverse set of target and efflux mutations (Table 1). From each strain three parallel populations (A, B and C) and their corresponding WTs were used to initiate a total of 30 populations which we subjected to experimental evolution for 300 generations in the absence of antimicrobial selection (CIP-R_evolved_ and WT_evolved_). Susceptibility testing (IC_90_ measurements) following experimental evolution revealed that the initial collateral effects (Figure 1 A) displayed by *E. coli* strains were largely lost (Figure 1 B) including conserved patterns of collateral sensitivity towards gentamicin and fosfomycin that were observed in four of five genetic backgrounds and the widespread efflux mediated cross resistance patterns (4). At the same time ciprofloxacin resistance levels were reduced in populations founded by strains harboring combinations of *gyrA* and various efflux mutations, a phenomenon also observed by others (18). Given the strong linear correlation between fold changes in IC_90_ and minimal inhibitory concentration (MIC) (6)(19), as well as absolute number comparisons (4), we conclude that the levels of ciprofloxacin resistance were reduced to just below the clinical breakpoint of 0.5 mg/L (20). Notably, ciprofloxacin resistance levels were still 14 – 20-fold higher than ancestral WT IC_90_. On the other hand, populations founded by the ancestral strain K56-2 CIP-R, which only harbored drug target mutations, maintained resistance levels above the clinical breakpoint after experimental evolution (average IC_90_ = 7.55 mg/L). K56-2 CIP-R_evolved_ population A and K56-50 CIP-R_evolved_ population A both showed a marked increase of 17.3- and 21.3-fold in fosfomycin resistance after evolution. This is likely due to the complex mutational landscape caused by mutations in the DNA mismatch repair genes *mutS* and *mutL* in these populations (Supplementary table S1). We also observed a conserved sensitization towards trimethoprim-sulfamethoxazole (SXT). For population specific collateral effects and dose response curves see Supplementary figure S1 and Supplementary figure S2.

**Table 1:**
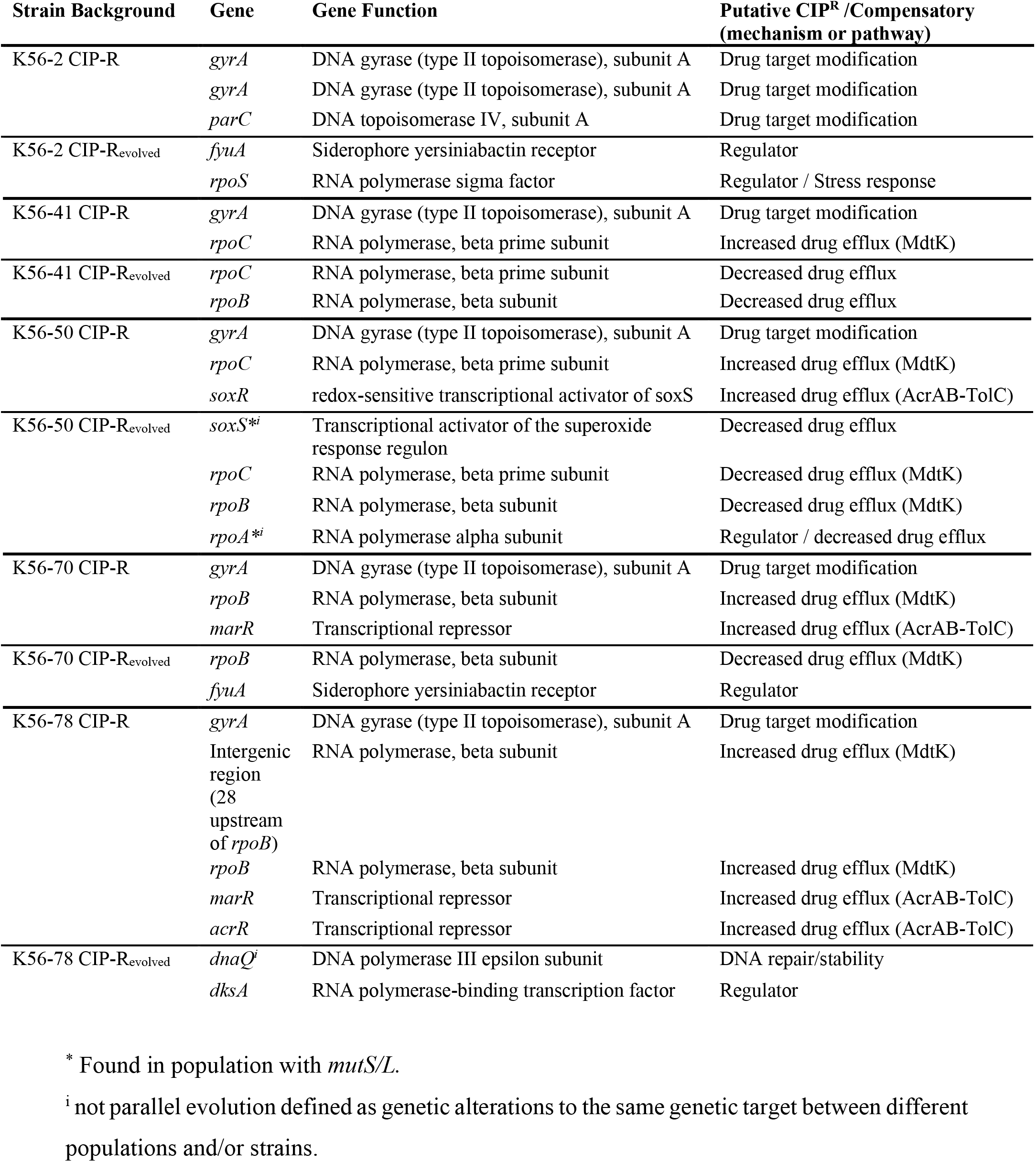
Genes with identified initial resistance mutations and putative compensatory mutations in clinical *E. coli* isolates after 300 generations in absence of antimicrobials.

| Strain Background | Gene | Gene Function | Putative CIP<sup>R</sup> /Compensatory (mechanism or pathway) |
| --- | --- | --- | --- |
| K56-2 CIP-R | *gyrA* | DNA gyrase (type II topoisomerase), subunit A | Drug target modification |
|  | *gyrA* | DNA gyrase (type II topoisomerase), subunit A | Drug target modification |
|  | *parC* | DNA topoisomerase IV, subunit A | Drug target modification |
| K56-2 CIP-R<sub>evolved</sub> | *fyuA* | Siderophore yersiniabactin receptor | Regulator |
|  | *rpoS* | RNA polymerase sigma factor | Regulator / Stress response |
| K56-41 CIP-R | *gyrA* | DNA gyrase (type II topoisomerase), subunit A | Drug target modification |
|  | *rpoC* | RNA polymerase, beta prime subunit | Increased drug efflux (MdtK) |
| K56-41 CIP-R<sub>evolved</sub> | *rpoC* | RNA polymerase, beta prime subunit | Decreased drug efflux |
|  | *rpoB* | RNA polymerase, beta subunit | Decreased drug efflux |
| K56-50 CIP-R | *gyrA* | DNA gyrase (type II topoisomerase), subunit A | Drug target modification |
|  | *rpoC* | RNA polymerase, beta prime subunit | Increased drug efflux (MdtK) |
|  | *soxR* | redox-sensitive transcriptional activator of *soxS* | Increased drug efflux (AcrAB-TolC) |
| K56-50 CIP-R<sub>evolved</sub> | *soxS*<sup>*i</sup> | Transcriptional activator of the superoxide response regulon | Decreased drug efflux |
|  | *rpoC* | RNA polymerase, beta prime subunit | Decreased drug efflux (MdtK) |
|  | *rpoB* | RNA polymerase, beta subunit | Decreased drug efflux (MdtK) |
|  | *rpoA*<sup>*i</sup> | RNA polymerase alpha subunit | Regulator / decreased drug efflux |
| K56-70 CIP-R | *gyrA* | DNA gyrase (type II topoisomerase), subunit A | Drug target modification |
|  | *rpoB* | RNA polymerase, beta subunit | Increased drug efflux (MdtK) |
|  | *marR* | Transcriptional repressor | Increased drug efflux (AcrAB-TolC) |
| K56-70 CIP-R<sub>evolved</sub> | *rpoB* | RNA polymerase, beta subunit | Decreased drug efflux (MdtK) |
|  | *fyuA* | Siderophore yersiniabactin receptor | Regulator |
| K56-78 CIP-R | *gyrA* | DNA gyrase (type II topoisomerase), subunit A | Drug target modification |
|  | Intergenic region (28 upstream of *rpoB*) | RNA polymerase, beta subunit | Increased drug efflux (MdtK) |
|  | *rpoB* | RNA polymerase, beta subunit | Increased drug efflux (MdtK) |
|  | *marR* | Transcriptional repressor | Increased drug efflux (AcrAB-TolC) |
|  | *acrR* | Transcriptional repressor | Increased drug efflux (AcrAB-TolC) |
| K56-78 CIP-R<sub>evolved</sub> | *dnaQ*<sup>i</sup> | DNA polymerase III epsilon subunit | DNA repair/stability |
|  | *dksA* | RNA polymerase-binding transcription factor | Regulator |

\* Found in population with *mutS/L*.
<sup>i</sup> not parallel evolution defined as genetic alterations to the same genetic target between different populations and/or strains.

**Figure 1:**
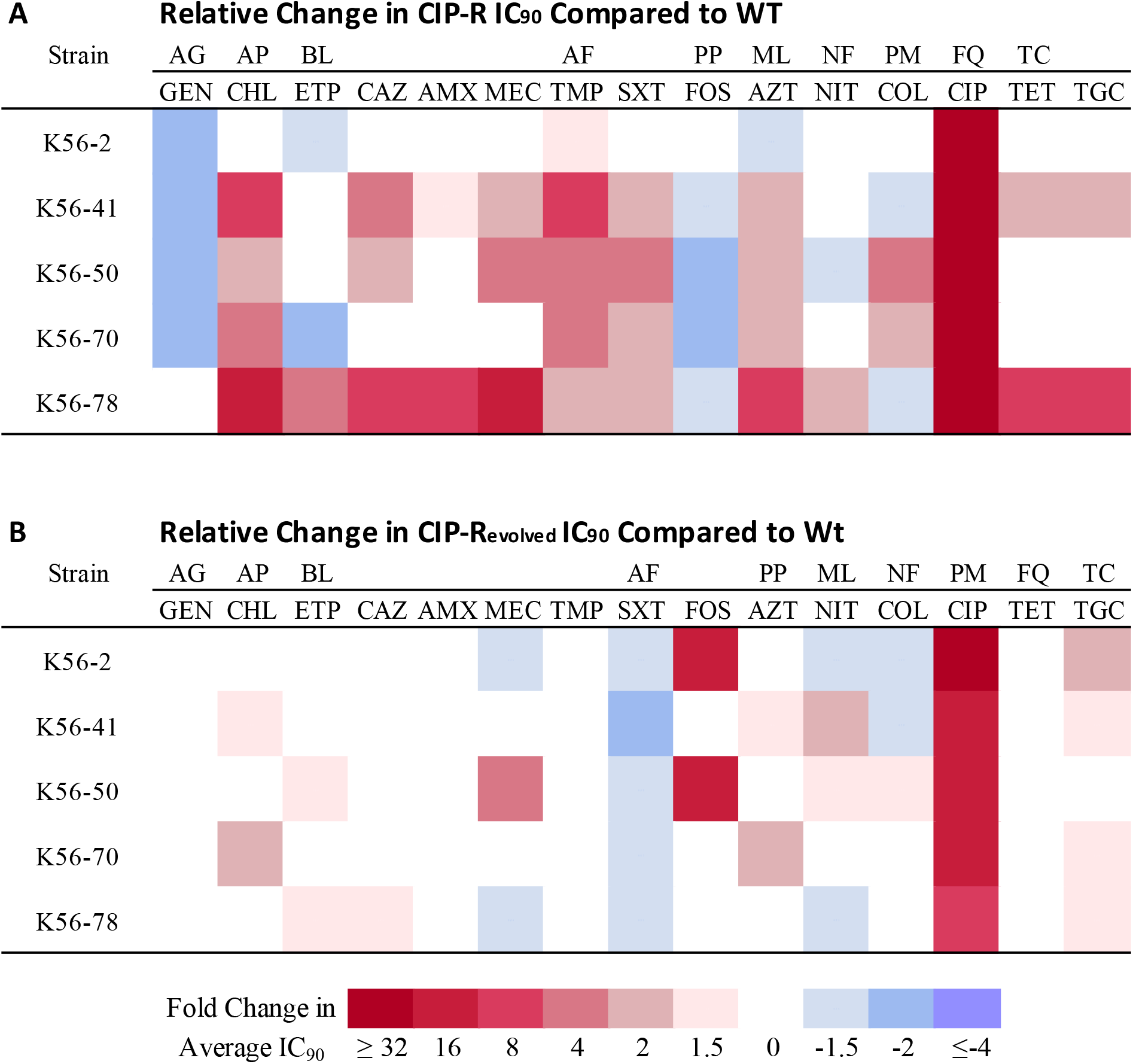
Collapsed collateral networks in ciprofloxacin resistant mutants following experimental evolution (300 generations) in the absence of antimicrobial selection. (**A**) Heatmap showing fold-change difference in susceptibility of ciprofloxacin resistant mutants compared to their respective WT tested towards a panel of 15 antimicrobials (Data previously reported in (4)). (**B**) Heatmap showing fold-change difference in susceptibility of ciprofloxacin resistant mutants compared to WT after 300 generation in absence of antimicrobial selective pressure. The reported values (Figure 1B) are based on pooled data from the three different parallel evolved populations for each strain. Drug classes: Aminoglycoside (AG), Amphenicol (AP), Beta-lactam (BL), Antifolate (AF), Phosphonic (PP), Macrolide (ML), Nitrofuran (NF), Polymyxin (PM), Fluoroquinolone (FQ) and Tetracycline (TC). Antimicrobials from left to right: Gentamicin (GEN), Chloramphenicol (CHL), Ertapenem (ETP), Ceftazidime (CAZ), Amoxicillin (AMX), Mecillinam (MEC), Trimethoprim (TMP), Trimethoprim-Sulfamethoxazole (SXT), Fosfomycin (FOS), Azithromycin (AZT), Nitrofurantoin (NIT), Colistin (COL), Ciprofloxacin (CIP), Tetracycline (TET) and Tigecycline (TGC).

### Collapse of collateral networks is linked to compensatory evolution

We previously reported that the fitness cost of resistance was a principal contributor to collateral effects (4). In this study, we hypothesized that the observed evolutionary instability of collateral effects (Figure 1) was driven by compensatory evolution. To test this hypothesis, we used relative growth rates as a proxy for fitness before and after experimental evolution. All evolved ciprofloxacin resistant mutants displayed significant ameliorations of initial fitness costs relative to WT controls (all p < 0.05, Figure 2), which were also evolved for 300 generations to control for putative effects of media adaptations. In strains harboring combinations of *gyrA* and various efflux mutations, relative fitness was fully restored (all p > 0.97, Figure 2, Supplementary table S2), except for K56-78 CIP-R_evolved_ (p < 0.0001; Figure 2, Supplementary table S2). K56-78 CIP-R had the most complex resistance mutation profile affecting two different efflux pumps (MdtK and AcrAB-TolC) in addition to *gyrA* (Table 1).

**Figure 2:**
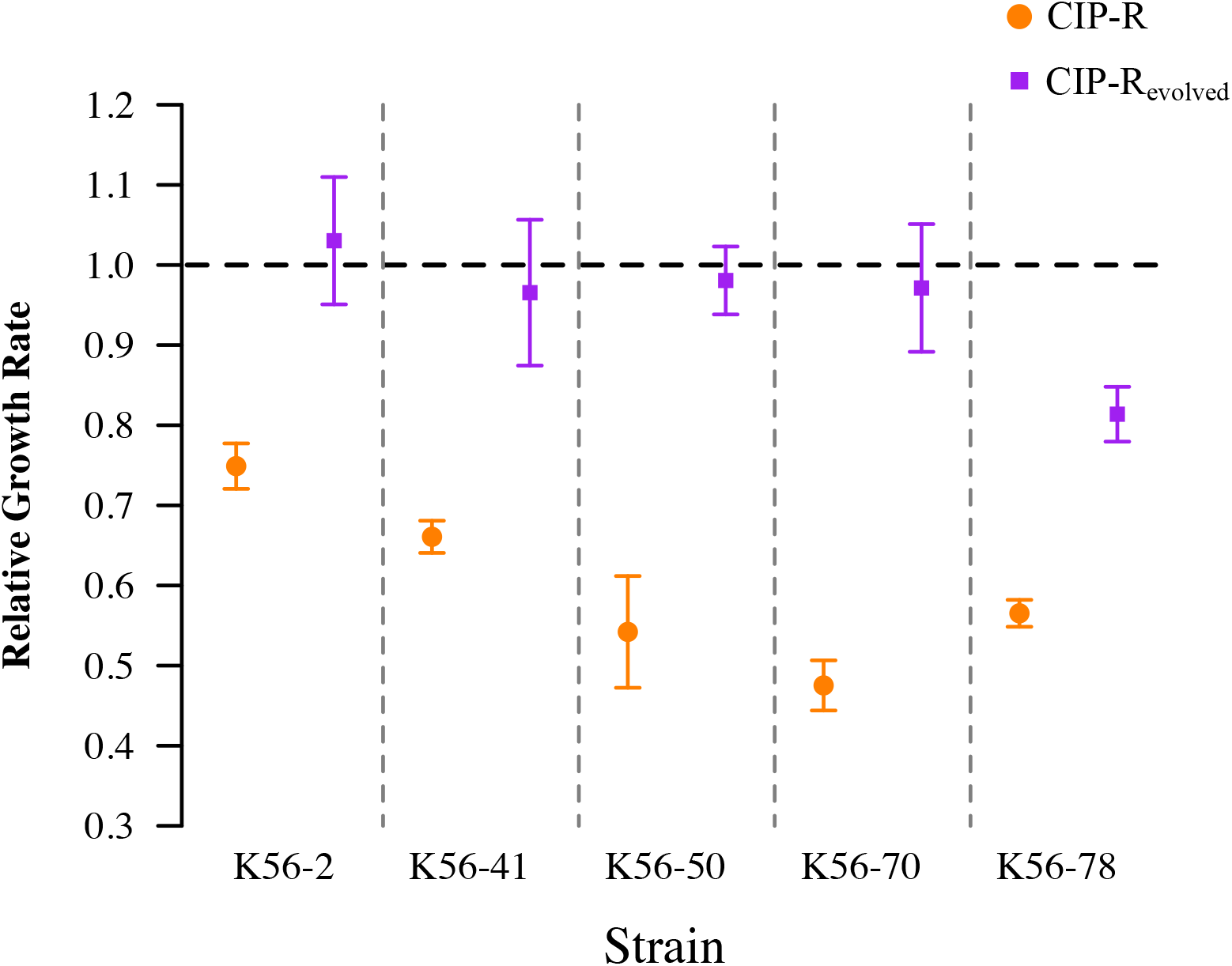
Relative growth rate of ciprofloxacin resistant mutants compared to their respective WT before* (orange) and after (purple) experimental evolution. Evolved ciprofloxacin resistant populations were compared to evolved WT populations to ameliorate fitness effects caused by media adaptation. Values below 1 (horizontal dashed line) denote a decreased growth rate in resistant mutants compared to their respective WT. We observed a reduction in the cost of ciprofloxacin resistance after evolution in all tested strains (p < 0.05). However, K56-78 CIP-R_evolved_ was the only strain that still displayed a significantly reduced growth rate compared to WT after evolution (P < 0.0001). Error bars denote the 95% confidence interval, n = 9. *Data previously reported in (4)

### Patterns of strong parallel evolution suggest compensatory mutations

To identify the genetic basis for the observed changes in collateral effects as well as potential compensatory mutations we subjected one fitness representative isolate from each of the evolved populations to whole genome sequencing (n=30; 1 clone from each CIP-R_evolved_ and WT_evolved_ population). After evolution, all CIP-R_evolved_ strains displayed putative compensatory mutations as indicated by mutations in genes modulating antimicrobial resistance, by parallel evolution, or both (Table 1). Parallel evolution is in general regarded as strong evidence for compensatory mutations (10)(21)(22).

We observed parallel evolution for two separate mutations in *fyuA* (F87L popB; Q507P pop C) as well as an identical *rpoS* (T298I) mutation in these two separate populations of K56-2 CIP-R_evolved_. FyuA is a TonB dependent yersinabactin siderophore receptor associated with uropathogenic *E. coli* (23)(24). A mutation in *fyuA* (Q516P) was also identified in K56-70 CIP-R_evolved_. RpoS is a sigma factor subunit of the *E. coli* RNA polymerase and acts as regulator in several cellular mechanisms (25)(26). The RpoS interaction with the SOS response has also been linked to maintenance of genome stability, protection from genotoxic stressors, altered susceptibility towards ciprofloxacin (27)(28) and as a target for compensatory mutation in fluoroquinolone resistant *Shigella sonnei* (29). It has previously been shown that *gyrA* mutations reduce DNA supercoiling and cause upregulation of RpoS in *Salmonella enterica* (30). Moreover, different *gyrA* mutations have also been shown to reduce DNA supercoiling in *E. coli* to a variable degree (31), which in turn has been associated with regulation of TonB gene expression (32)(33). Strong parallel evolution in our experimental evolution assay combined with existing literature indicate that RpoS and potentially FyuA as a TonB dependent receptor could function as targets for compensatory mutations.

In addition to the drug target mutation *gyrA* (S83L) in K56-41 CIP-R, K56-50 CIP-R, K56-70 CIP-R and K56-78 CIP-R, all strains displayed mutations in genes related to drug efflux (4). Following experimental evolution, these strains also acquired additional mutations hypothesized to restore efflux activity to WT levels. K56-41 CIP-R initially had a 9 bp deletion in *rpoC* and mutations in this gene have been associated with increased activity of the MdtK efflux pump (34). Following experimental evolution we identified additional mutations in *rpoC* as well as in *rpoB*, which are also known to affect the MdtK efflux pump (34). These results indicate that the MdtK efflux system is the primary target for compensatory mutations in this strain background. In K56-50 CIP-R_evolved_ we identified a conversion of the initial RpoC S86F mutation into a F86C mutation in two of the three evolved populations (B and C), which likely restored WT RpoC activity due to the more similar physiochemical properties of cysteine and serine (WT). Population C also had an additional mutation in *rpoB* (V857E). K56-70 CIP-R displayed an initial mutation in RpoB (I668F) and a deletion in *marR*, respectively. All of the evolved K56-70 CIP-R_evolved_ populations displayed amino acid changes in the same location as the initial amino acid change in RpoB (F668S, F668V and F668L) indicating that the initial RpoB (I668F) mutation is a highly selective target for compensatory reversion – mutations. Strain K56-78 CIP-R displayed mutations affecting both MdtK and the AcrAB-TolC efflux pumps. After experimental evolution, K56-78 CIP-R_evolved_ did not fully restore cost of resistance and parallel evolution was observed only in *dksA* in two of the evolved populations. DksA has previously been associated with ciprofloxacin tolerance functioning as a regulator of RNA polymerase in *Yersinia pseudotuberculosis* (35) as well as part of the ppGpp regulation in *E. coli* (36). If, and how these mutations affect efflux levels is currently unknown. In two of the evolved CIP-R populations (K56-2 CIP-R_evolved_ pop A and K56-50 CIP-R_evolved_ pop A) and in two of the evolved WT populations (K56-41_evolved_ pop B and K56-78_evolved_ pop C) we identified mutations in *mutS* or *mutL*. Although this suggests a potential adaptive benefit of increased mutation rates in our evolution assay, these populations were handled with caution in the downstream analysis due to their complex mutational landscapes, Supplementary table S1.

### The cost of ciprofloxacin resistance mutations is dependent on genetic background

The cost of drug target mutations observed in our clinical isolate (K56-2 CIP-R) was markedly larger than in several earlier studies using emblematic laboratory strains of *E. coli* (37)(38), although variation in literature exists with reports of larger costs more similar to our results (39). To further investigate the cost of these mutations in a different genetic background, we inserted the three initial mutations found in K56-2 CIP-R (*gyrA* S83L; *gyrA* A119E; *parC* G78D) into *E. coli* MG1655. In the *E. coli* MG1655 background our results were similar to earlier studies (37)(38), w = 0.95 (standard deviation = 0.031, n=3). No other mutations were observed in K56-2 CIP-R (4) that could explain the difference in fitness cost and we conclude that the large fitness effects observed here are due to strain specific differences, *i.e*. genetic background.

### Ancestral efflux phenotype was restored after evolution in absence of antimicrobials

To verify that initial ciprofloxacin resistance was due to the predicted increased efflux before experimental evolution, we functionally characterized the four strains with combinations of *gyrA* and efflux mutations (4). The expression of five structural efflux pump genes from efflux pumps (AcrAB-TolC (40), MdtK (NorE) (34) and MdfA (41)) with ciprofloxacin as one of their substrates were assessed by reverse transcriptase quantitative PCR (RT-qPCR). All four strains displayed increased transcription of *mdtK* and at least two genes associated with the AcrAB-TolC efflux pump compared with WT (Figure 3). In addition, we observed increased expression of *mdfA* in all strains except K56-78 CIP-R. We recently demonstrated that the majority of collateral responses in these strains were due to mutations in efflux regulatory genes and not in *gyrA* (4). We further hypothesized that the loss of collateral phenotypes following experimental evolution was due to restored efflux pump activity in the absence of antimicrobial selection. RT-qPCR on evolved strains verified significant reduction in expression of various efflux systems across all tested strains, strongly suggesting that reduced efflux levels were responsible for loss of collateral responses as well as mitigations of fitness costs. The incomplete restoration of efflux systems to WT K56-78 CIP-R also provides an explanation for the observed partial fitness mitigation compared to the other strains (Figure 3). The functional assessment of efflux genes tested here suggest that restoration of WT efflux phenotype acts as substrate/driver for fitness compensatory evolution and as primary sources for loss of collateral sensitivity and resistance networks. Many of the antimicrobials used in this study are known substrates of the efflux pumps investigated in this study (Table 2).

**Figure 3:**
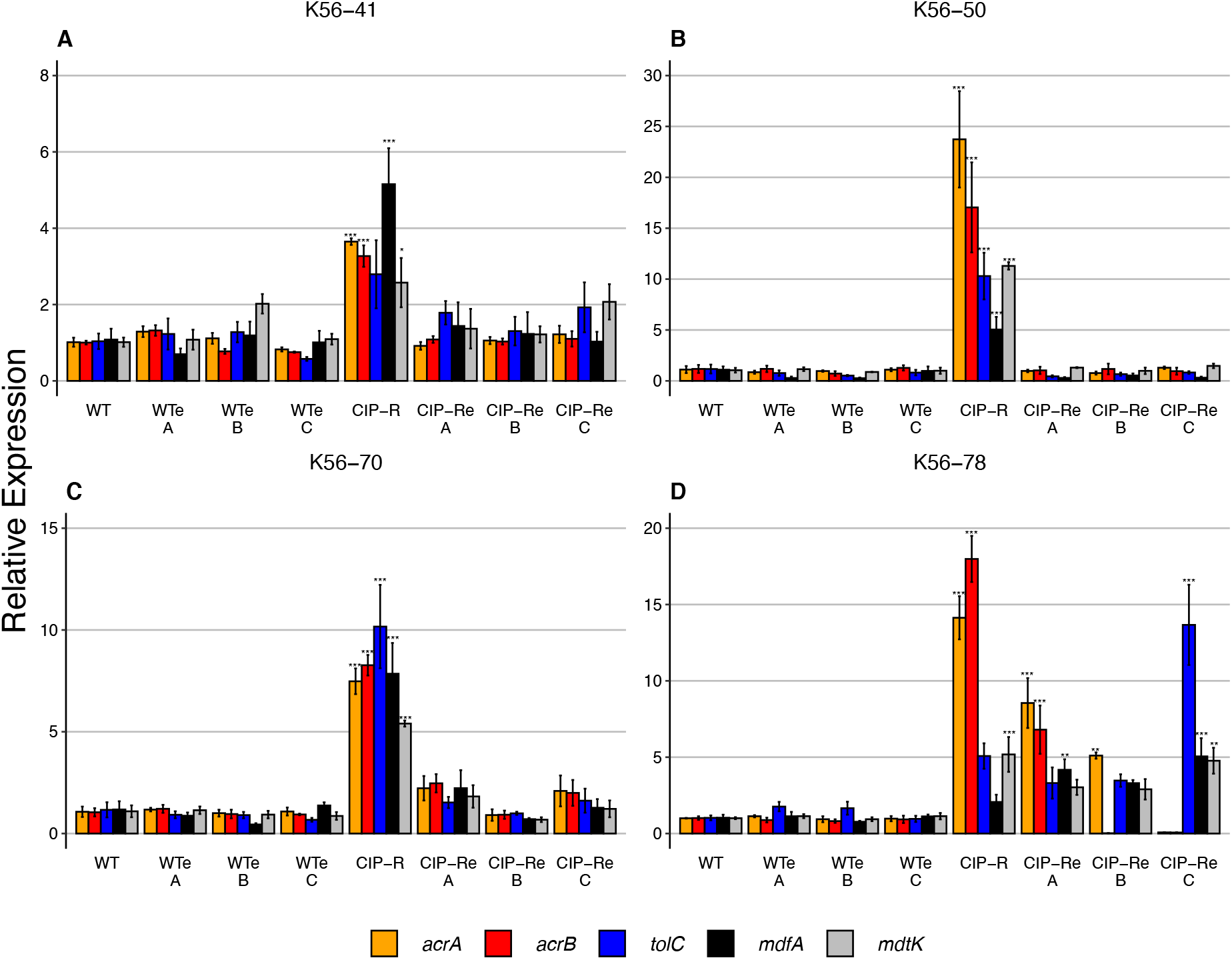
Relative transcription levels of identified efflux genes compared to the corresponding WT in the CIP-R mutants and evolved WT_evolved_ (WT_e_) and CIP-R_evolved_ (CIP-R_e_) mutants using reverse transcriptase quantitative PCR. A, B and C represent the different evolved populations for each strain. Increased efflux expression in CIP-R strains were restored to WT levels following evolution in all strains but K56-78. Whiskers denote standard error, significance (***0.0001**0.001,*0.05) from WT expression was adjusted for multiple comparisons with Dunnet’s test, n = 3.

**Table 2:** Drugs or drug classes used in this study known to interact with identified and related efflux pumps.

| Efflux Pump | Antimicrobials |  |  |  |  |  |  |  |  |  |  |  |  |  | References |
| --- | --- | --- | --- | --- | --- | --- | --- | --- | --- | --- | --- | --- | --- | --- | --- |
|  | AMX | MEC | CAZ | ETP | FOS | COL | TET | TGC | GEN | AZT | CHL | CIP | NIT | SXT |  |
| AcrAB-TolC | ◇ | ◇ | ◇ | ◇ |  |  | + | ◇ |  | ◇ | + | + |  | + | (40)(42)(43) |
| MdtK | ◇ | ◇ | ◇ | ◇ | + |  |  |  |  |  | + | + |  | ◇ | (34)(44) |
| MdfA |  |  |  |  |  |  | + | ◇ | ◇ | ◇ | + | + |  | ◇ | (41)(44) |
| *Other efflux pumps known to require AcrA and TolC* |  |  |  |  |  |  |  |  |  |  |  |  |  |  |  |
| AcrAD-TolC | ◇ | ◇ | ◇ | ◇ |  |  |  |  | + |  |  | ◇ |  |  | (43)(44)(45) |
| *Other relevant efflux pumps known to require TolC* |  |  |  |  |  |  |  |  |  |  |  |  |  |  |  |
| AcrEF-TolC | ◇ | ◇ | ◇ | ◇ |  |  | + | ◇ |  | ◇ | + | + |  | ◇ | (42)(43)(44)(46) |
| MdtABC-TolC | ◇ | ◇ | ◇ | ◇ |  |  |  |  |  |  |  | ◇ |  |  | (43)(45)(47)(48) |
| MdtEF-TolC | ◇ | ◇ | ◇ | ◇ |  |  |  |  |  | ◇ |  | + |  |  | (43)(49) |
| EmrAB-TolC |  |  |  |  |  |  |  |  |  |  |  | ◇ |  |  |  |

+, evidence of efflux; ◇, evidence of drug class efflux.
Antimicrobials from left to right: Amoxicillin (AMX), Mecillinam (MEC), Ceftazidime (CAZ), Ertapenem (ETP), Fosfomycin (FOS), Colistin (COL), Tetracycline (TET), Tigecycline (TGC), Gentamicin (GEN), Azithromycin (AZT), Chloramphenicol (CHL), Ciprofloxacin (CIP), Nitrofurantoin (NIT) and Trimethoprim-Sulfamethoxazole (SXT).

## Discussion

Clinical use of collateral networks to reduce emergence and spread of antimicrobial resistance is contingent on a robust identification and predictions of collateral responses informed by susceptibility testing as well as rapid detection of resistance mechanisms (4). Here we demonstrate evolutionary instability of collateral networks in ciprofloxacin resistant clinical *E. coli* UTI strains following 300 generations of experimental evolution in the absence of antimicrobial selective pressures. Our data strongly suggest that the evolutionary instability is caused by compensatory mutations reducing the initial fitness costs of ciprofloxacin resistance. In a recent study by Barbosa *et al*. they investigated evolutionary stability of collateral sensitivity from a different perspective, subjecting *P. aeruginosa* to sequential exposure of antimicrobial pairs for which reciprocal collateral sensitivity was demonstrated (3). They identified several cases of re-sensitization to the first antimicrobial in a switch protocol following resistance development towards the second drug. This bacterial response occurred more frequently than development of multi-drug resistance demonstrating that reciprocal collateral sensitivity is promising from a treatment perspective and effectively restrict bacteria in a “double bind evolutionary trap” (3). Our study is highly relevant downstream from the results presented by Barbosa *et al*. and underscore the importance of considering temporal evolutionary dynamics when predicting collateral networks.

Efflux mediated ciprofloxacin resistance, the predominant resistance factor observed in the clinical isolates tested here, is widespread in the clinical setting, and has been described in several reports investigating collateral networks across multiple species (9) (15). The high prevalence of resistance caused by altered efflux phenotype is likely due to the large mutational target and high selection pressure protocols used to select for resistant isolates for comparison with selective ancestors. Although efflux is of clinical relevance in *E. coli* it is not as prevalent as drug target mutations (50)(51)(52)(53). However, here we also identified instability of collateral effects in a mutant containing only drug target mutations suggesting a generality of our findings with respect to ciprofloxacin resistant *E. coli*. This result suggests that, at least for newly acquired resistance determinants, collateral networks should be treated as temporally unstable. The observation that fitness costs of resistance is a principal contributor of collateral networks (4) may not be universal and is likely linked to substantial initial fitness costs of newly acquired resistance determinants. This is supported by reports using *E. coli* MG1655 (2) and *Streptococcus pneumoniae* (54) where this link was not observed. Taken together with a recent report demonstrating that collateral networks are dependent on environmental factors (17), our data support that collateral responses measured for combinations of host strains and newly acquired resistance determinants across non-standardized growth conditions serve as poor references for accurate predictions. To that end, future clinical application of collateral responses requires careful considerations of evolutionary stability, impact of biotic and abiotic factors on quantification and predictions of collateral effects as well as pharmacodynamics and pharmacokinetics of drug-bacteria combinations.

## Methods

### Bacterial strains and strain constructions

In this study we assess five clinical isolates of urinary tract infection *E. coli* from the ECO-SENS collection (55) that were previously selected for resistance *in vitro* (Table 1) by plating on increasing concentrations of ciprofloxacin (4). *E. coli* ATCC 25922 was used as a control strain in all IC_90_ assays. *E. coli* MG1655 was used as the recipient for *gyrA* (S83L; A119E) and *parC* (G78D) drug target mutations following the pORTMAGE protocol (56) (57) using the pORTMAGE-2 plasmid as described in (57). Oligonucleotides introducing the three different mutations used in the pORTMAGE cycling protocol was designed using MODEST (58). See Supplementary table S3 and S4 for complete list of bacterial strains and primers used in this study.

### Evolution in absence of antimicrobials

From five ciprofloxacin resistant (CIP-R) mutants (4) three parallel populations (A, B and C) and their corresponding WTs resulting in a total of 30 populations were evolved for 300 generations in absence of antimicrobial pressure using a 1:100 serial transfer assay regime resulting in approximately 6.64 generations per transfer. Utilizing 96 deep-well plates, 10 μl from stationary phase cultures were transferred into 990 μl fresh Muller-Hinton II broth (MHB; Becton, Dickinson and Company (BD), Franklin Lakes, NJ, USA) every 24 hours and incubated at 37 °C shaking at 500 rpm. Evolving populations were grown in a checkerboard pattern with un-inoculated growth medium to control for absence of cross-contamination.

### Susceptibility networks

Susceptibility networks towards 15 antimicrobials (see Figure 1) were obtained by assessing change in susceptibility testing using a broth microdilution assay (IC_90_ determination) (4)(2) before and after evolution. Overnight cultures of bacteria were suspended in sterile saline to make a 0.5 McFarland standard and diluted 1:1000 in MHB. 100 μL were transferred into 96-well microtiter plates containing serial dilution of one of the 15 antibiotics following 1.5-fold dilution steps; 200 μL final volume. The plates were incubated shaking at 37°C, 700 rpm for 18 hours. OD_600_ was measured in a Versamax plate reader (Molecular Devices Corporation, CA, USA). IC_90_ was calculated as OD_600_ antibiotic concentration * OD_600_ positive control^-1^ (2). Results are based on at least three biological replicates performed on different days and all plates included *E. coli* ATCC 25922 test strain as control.

### Growth Rates

Growth curves of the evolved populations of WT_evolved_ and CIP-R_evolved_ was obtained by inoculating at three biological replicates into 2 mL of MHB which was incubated at 37 °C, 500 rpm for 24 hours. The starting cultures were then diluted 1:100 resulting in starting titer of ~2 x 10^7^ cells mL^-1^. From these dilutions 250 μL was added to a 96-well microtiter plate in triplicates. The plate was incubated overnight in a Versamax plate reader (Molecular Devices Corporation, California, USA) at 37°C and shaking for 9.2 minutes between each read. Optical Density (OD_600_) measurements were taken every 10 minutes and growth rates (r) was estimated using GrowthRates v.2.1(59). The reported relative growth rates (w) were obtained using the formula r_(CIP-Revolved)_ x r_(WTevolved)_^-1^. One-way ANOVAs adjusted for multiple comparisons with Tukey HSD were used to assess significant changes in growth rates. Significance was considered p ≤ 0.05

### Whole genome sequencing

Genomic DNA was isolated from a single colony of a fitness representative biological replicate in the growth rates assay from each of the parallel evolved CIP-R and WT populations. We used GenElute for Bacteria Genomic DNA kit (Sigma-Aldrich, St. Louis, MO, USA) following the Gram-Positive DNA extraction protocol to increase DNA isolation yield in our clinical strains. Purity of the samples was assessed using NanoDrop (Thermo Fisher) and DNA quantification performed using a Qubit High Sensitivity DNA assay (Life Technologies). Sequencing was performed according to the Nextera XT DNA library prep kit (Illumina, San Diego, CA, USA) at the Genomics Resource Centre Tromsø (UiT Arctic University of Norway). Downstream analysis of DNA sequences was performed according to Podnecky *et al*. 2018 (4) to allow for direct comparisons of the sequence data. In short, raw reads from the evolved WT_evolved_ and CIP-R_evolved_ were aligned to previously annotated (RAST 2.0 for *E. coli* (60)) sequences of the WT strains utilizing standard settings in SeqMan NGen (DNASTAR, Madison, WI). Minimum values of reported SNPs were 10x coverage depth and 90% variant base calls. Reported SNPs and indels were manually inspected.

### Relative gene expression

To assess relative transcription levels, reverse transcriptase quantitative PCR (RT-qPCR) was performed. Samples were grown in MHB overnight at 37°C with shaking (200 RPM). 500 μL of mid-log phase (OD_600_nm = 0.45 - 0.7, strain-specific) cells were stabilized using RNA Protect Bacteria Reagent (QIAGEN, Hilden, Germany), and cell pellets stored at −80°C. Total RNA was extracted using the RNeasy Mini Kit (Qiagen). 1 μg of RNA was treated using DNA-free DNA removal kit (Invitrogen, Carlsbad, CA, USA) with 2x concentration of rDNase I, as recommended for rigorous DNase treatment. cDNA synthesis was performed on 4 μL of DNase-treated RNA using SuperScript^®^ III First-Strand Synthesis SuperMix for qRT-PCR (Invitrogen). The DNase treatment and cDNA synthesis protocols were upscaled as needed. RT-qPCR was performed using the PowerUp™ SYBR™ Green Master Mix (Thermo Fisher) on a 7300 Real Time PCR System and 7300 System SDS RQ Study Software (Applied Biosystems, Beverly Hills, CA, USA). Melt curve analyses were used to rule out secondary products and primer dimer formation. Samples were tested in technical triplicate and Ct values were averaged for analysis. CysG was used for data normalization; primer sequences are listed in Table 3 Relative expression to the respective WT or CIP-R isolate was calculated using the ΔΔCt method (61). Relative expression was assessed in at least biological triplicate; additional replicates were performed if relative expression varied more than four-fold between biological replicates and the outlier excluded, if applicable. Average relative expression and standard error were calculated, Dunnett’s-test controlling for multiple comparisons were used to assess significant changes in gene expression. Significance was considered p ≤ 0.05.

**Table 3:**
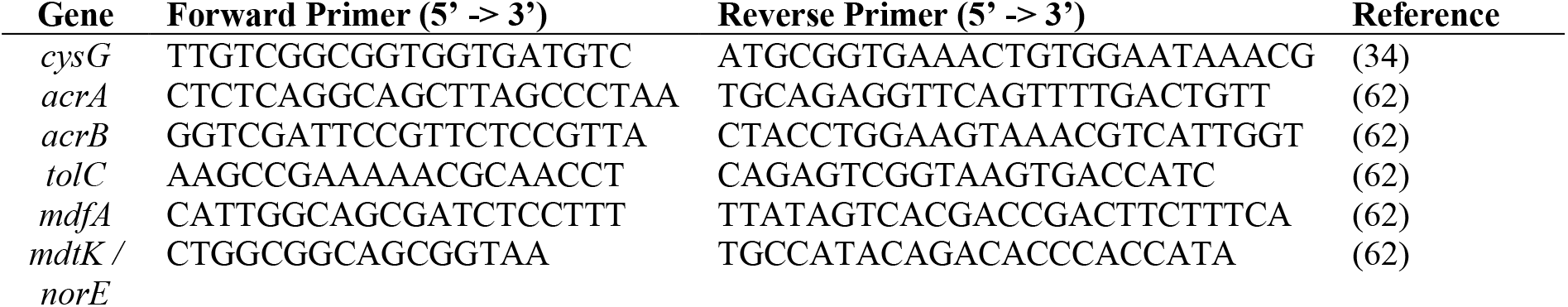
RT-qPCR primers used in this study.

### Data analysis and software

Statistical analyses were performed using R (version 4.0.3) (63) and GraphPad Prism (version 9.1.1), GraphPad Software, San Diego, California USA. Growth rates were obtained from GrowthRates version 2.1 (59).

## Acknowledgements

We would like to thank Hagar Taman and Prof. Ruth H. Paulssen at the UiT Genomic support Center Tromsø, for WGS Illumina sequencing service. We would also like to thank Christopher Frölich, João Gama, Francois Cleon, Jónína S. Gudmundsdóttir and Elizabeth G. A. Fredheim for valuable feedback, discussions and support.

## Author contribution

Conceptualization VS, PJJ, ØS. Methodology: VS, PJJ, NLP, KH. Investigation: VS, ELØ, AMS, NLP, KH. Formal analysis VS, NLP. Manuscript was written by VS, PJJ, NLP, ELØ and AMS with input from all other authors. Project administration and supervision: PJJ. Funding acquisition: PJJ.

## Data and materials availability

Data and material is available in the main text and supplementary files.

## Supplementary files

**Supplementary table S1 – Full list of ancestral and evolved mutations**

**Supplementary table S2 – Tukey HSD comparisons of growth rates**

**Supplementary table S3 – Full list of bacterial strains used in this study**

**Supplementary table S4 – Full list of primers and oligos used in this study**

**Supplementary figure S1 – Heatmap of population specific changes in IC_90_**

**Supplementary figure S2 – Dose response curves**

