## Supplementary figure S1 for "Evolutionary instability of collateral susceptibility networks in ciprofloxacin resistant clinical *Escherichia coli* strains"

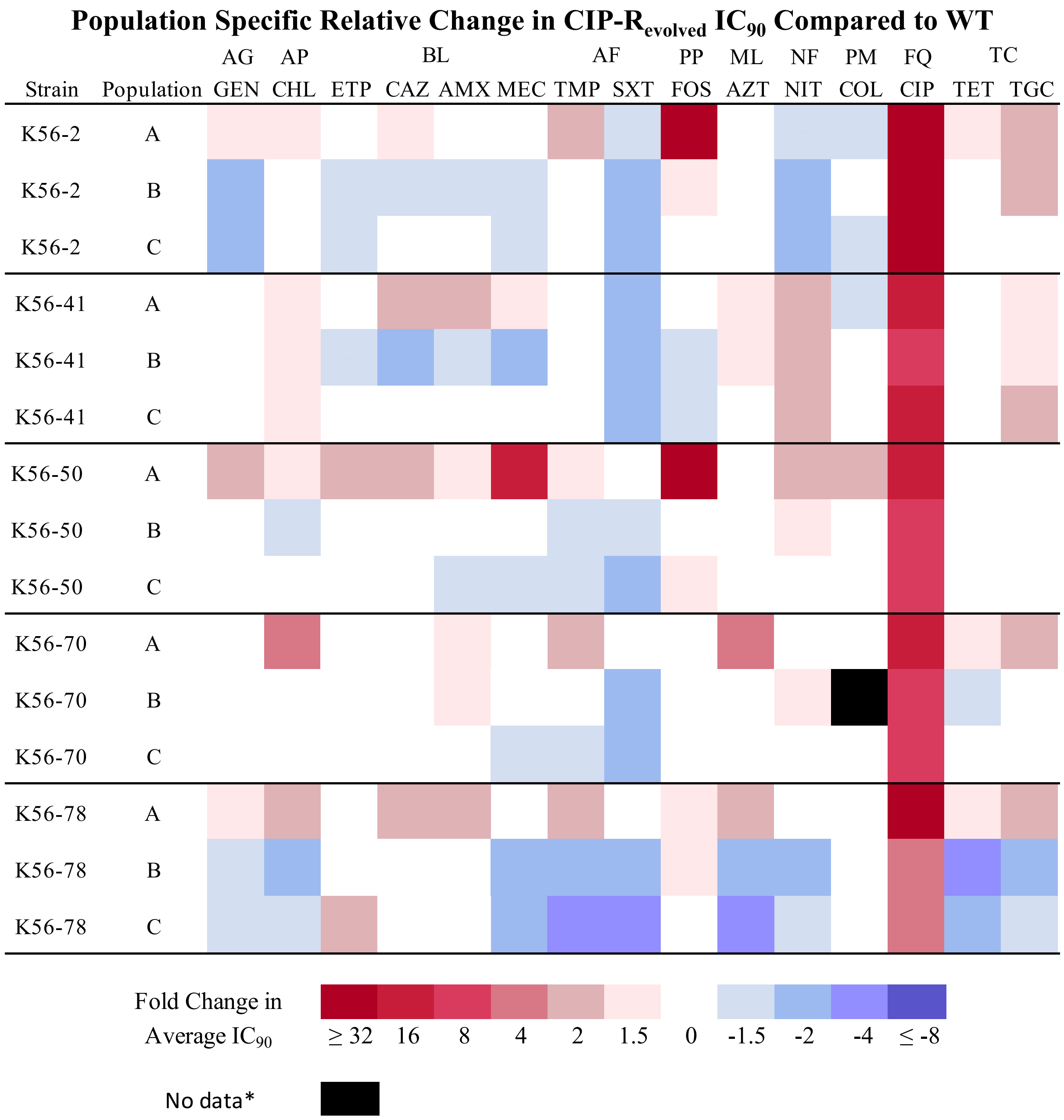


**Supplementary figure 1:** Heatmap showing population specific fold-change in susceptibility of ciprofloxacin resistant mutants compared to their corresponding ancestral WT after 300 generation in absence of antimicrobial selective pressure. Drug classes: Aminoglycoside (AG), Amphenicol (AP), Beta-lactam (BL), Antifolate (AF), Phosphonic (PP), Macrolide (ML), Nitrofuran (NF), Polymyxin (PM), Fluoroquinolone (FQ) and Tetracycline (TC). Antimicrobials from left to right: Gentamicin (GEN), Chloramphenicol (CHL), Ertapenem (ETP), Ceftazidime (CAZ), Amoxicillin (AMX), Mecillinam (MEC), Trimethoprim (TMP), Trimethoprim-Sulfamethoxazole (SXT), Fosfomycin (FOS), Azithromycin (AZT), Nitrofurantoin (NIT), Colistin (COL), Ciprofloxacin (CIP), Tetracycline (TET) and Tigecycline (TGC).
