## Supplementary figure S2 for "Evolutionary instability of collateral susceptibility networks in ciprofloxacin resistant clinical *Escherichia coli* strains"

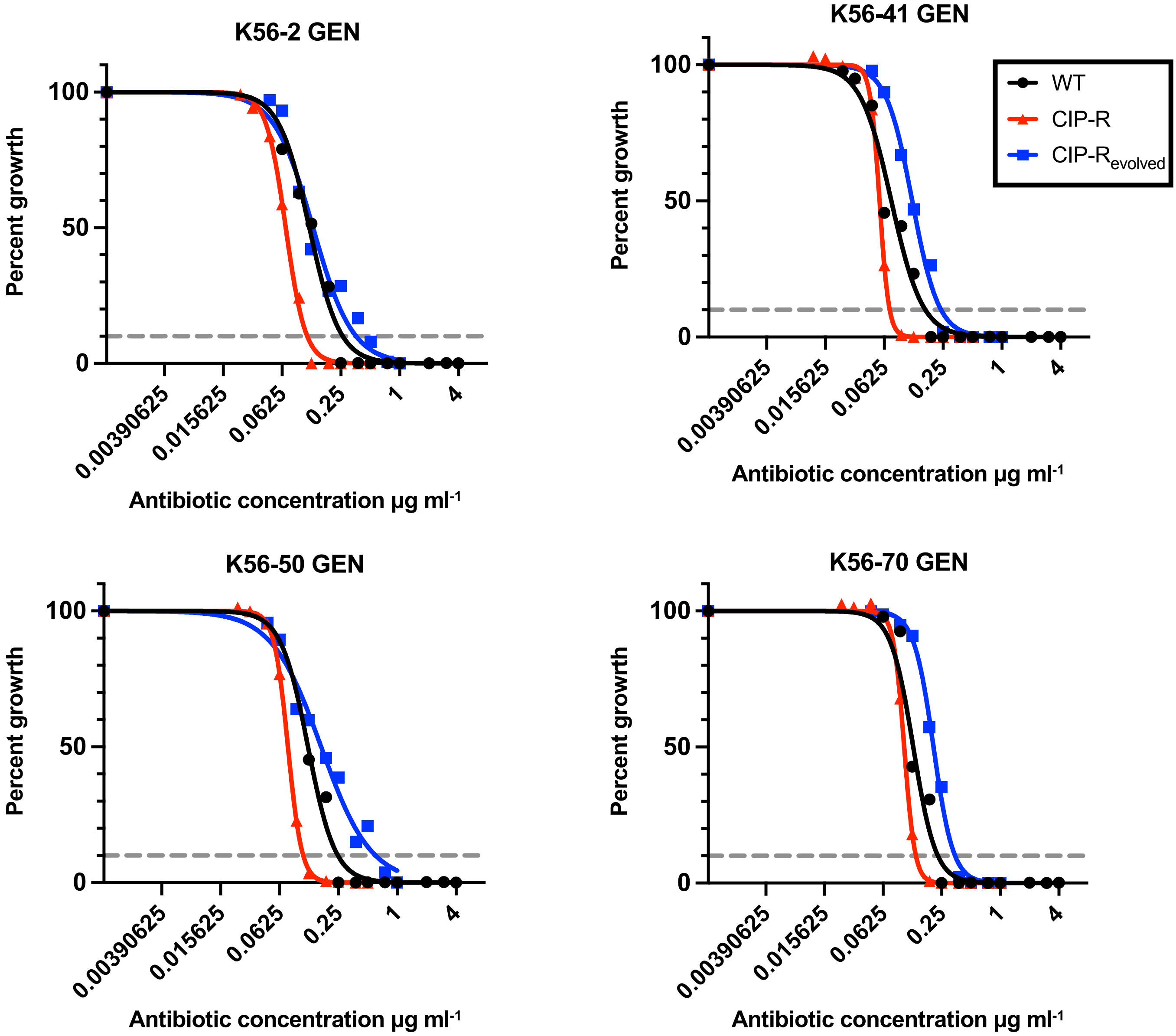


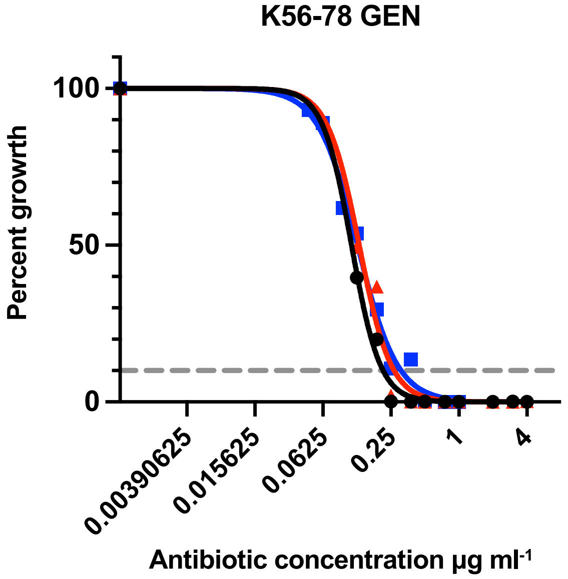


**Supplementary figure 2 – Gentamicin:**
Dose response curves denoting percent growth at given antibiotic concentrations for Ancestral WT (black circles), Ancestral ciprofloxacin resitant (red triangles) and Evolved ciprofloxacin restant (blue squares). Each strain background are separated into different graphs. Grey dashed line denotes the $IC_{90}$.

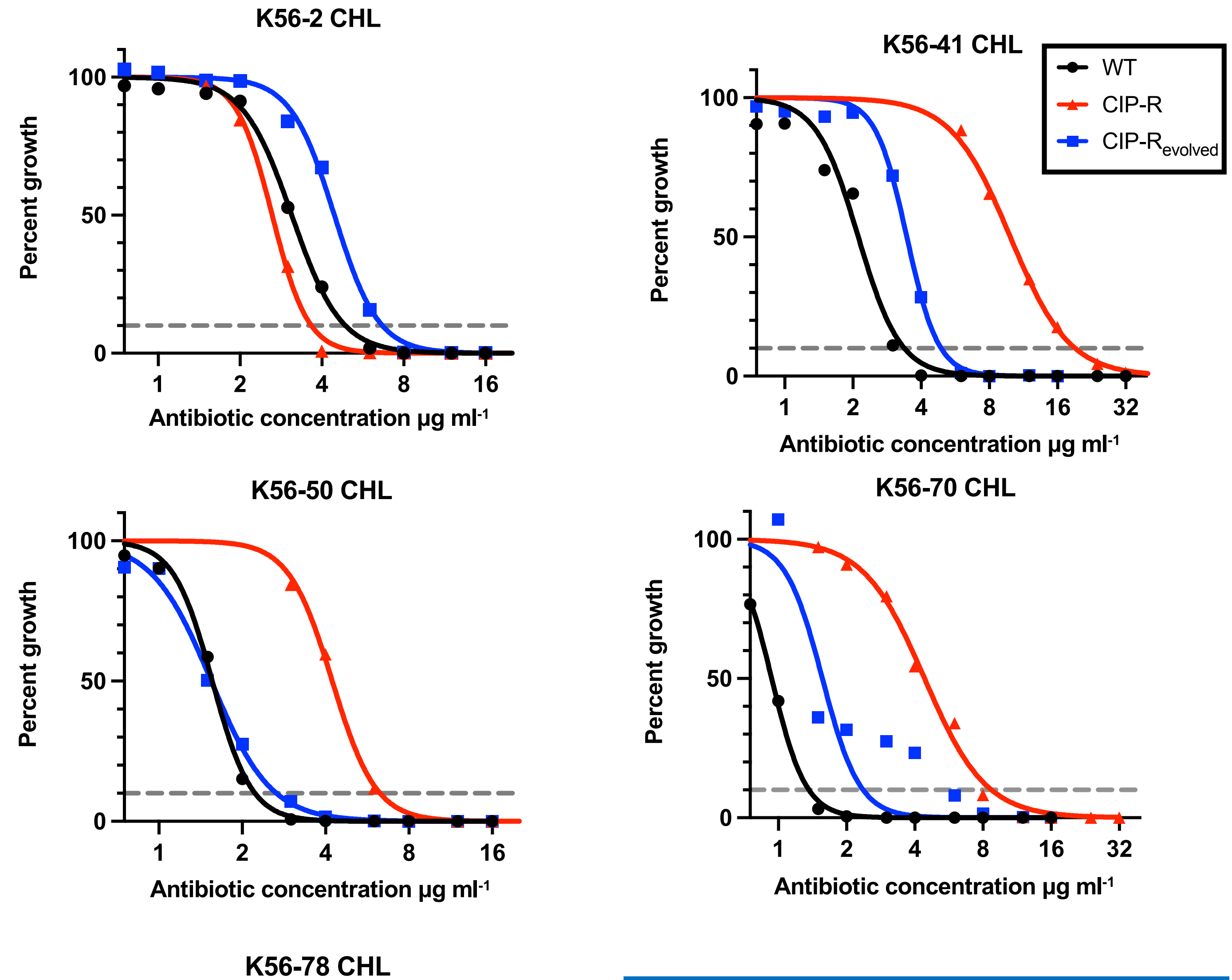


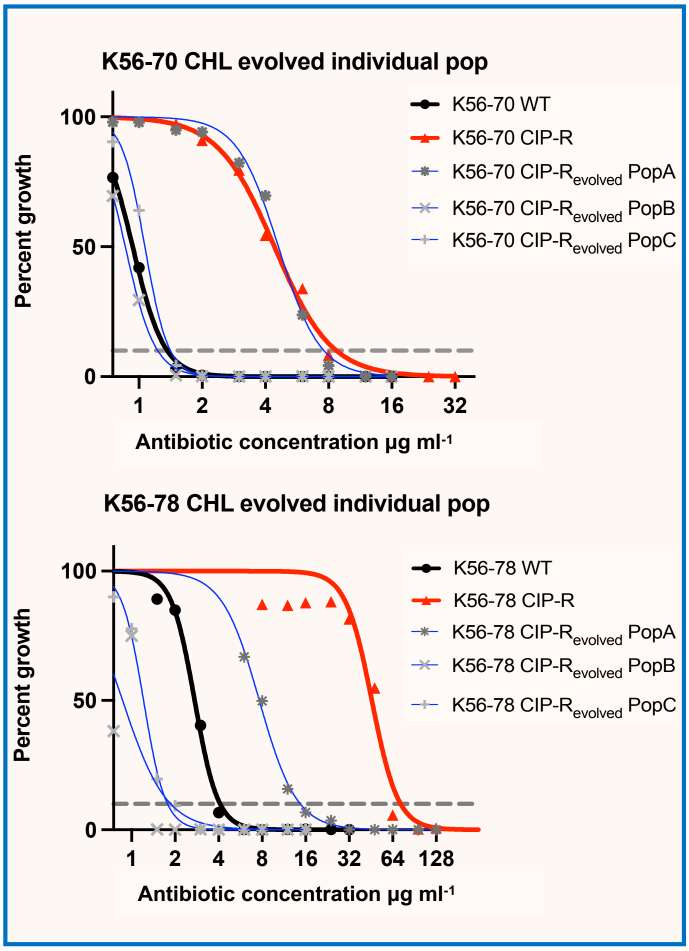


**Supplementary figure 2 – Chloramphenicol:**
Dose response curves denoting percent growth at given antibiotic concentrations for Ancestral WT (black circles), Ancestral ciprofloxacin resitant (red triangles) and Evolved ciprofloxacin restant (blue squares). Each strain background are separated into different graphs. Framed in the lower right corner the three individually evolved populations (grey symbols and blue lines) for strain K56-70 and K56-78 is shown separately due to large variation between the populations. Grey dashed line denotes the $IC_{90}$.

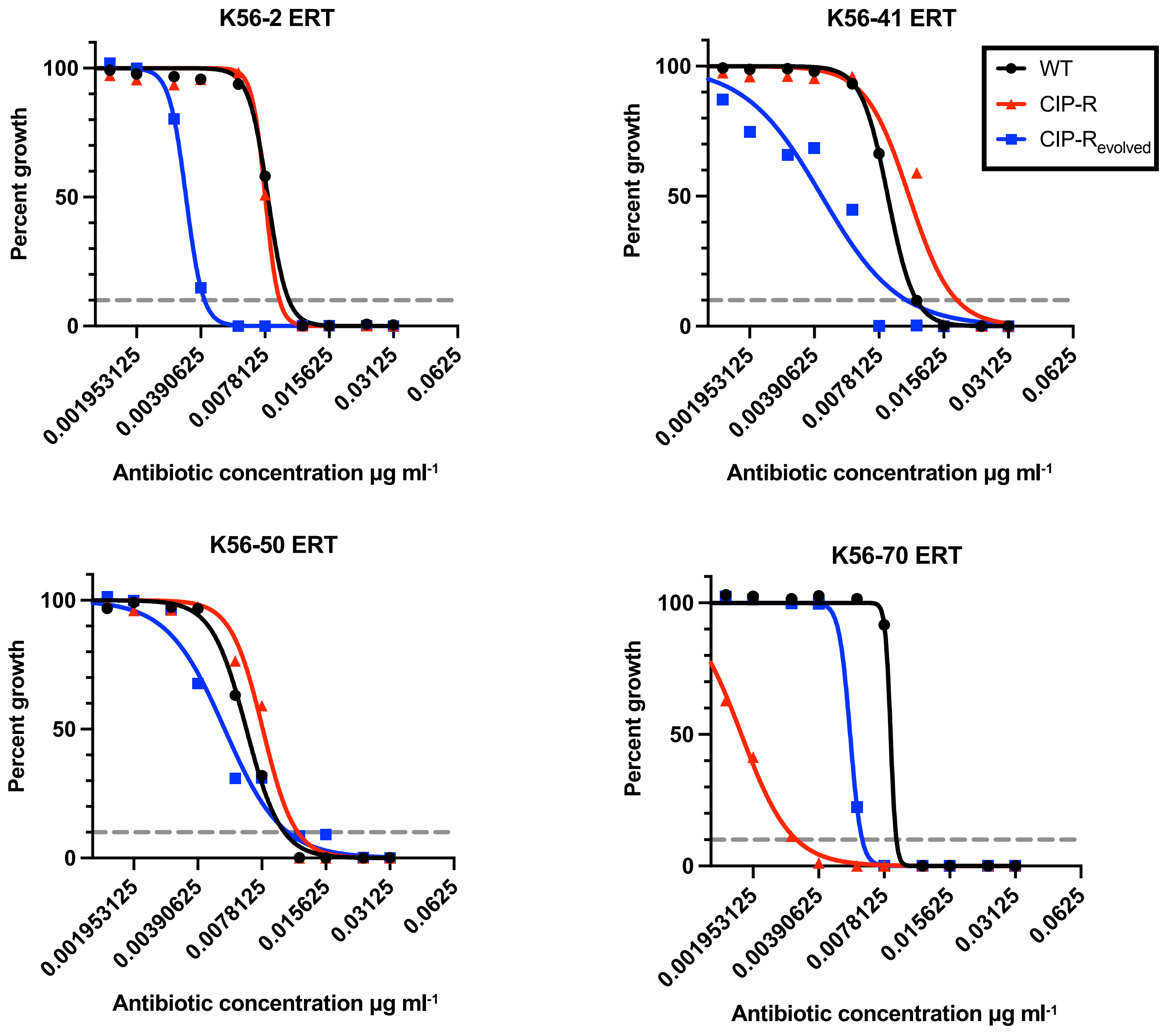


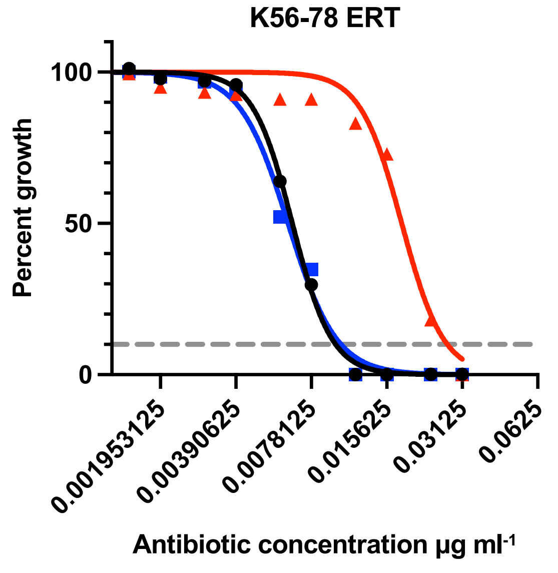


**Supplementary figure 2 – Ertapenem:**
Dose response curves denoting percent growth at given antibiotic concentrations for Ancestral WT (black circles), Ancestral ciprofloxacin resitant (red triangles) and Evolved ciprofloxacin restant (blue squares). Each strain background are separated into different graphs. Grey dashed line denotes the $IC_{90}$.

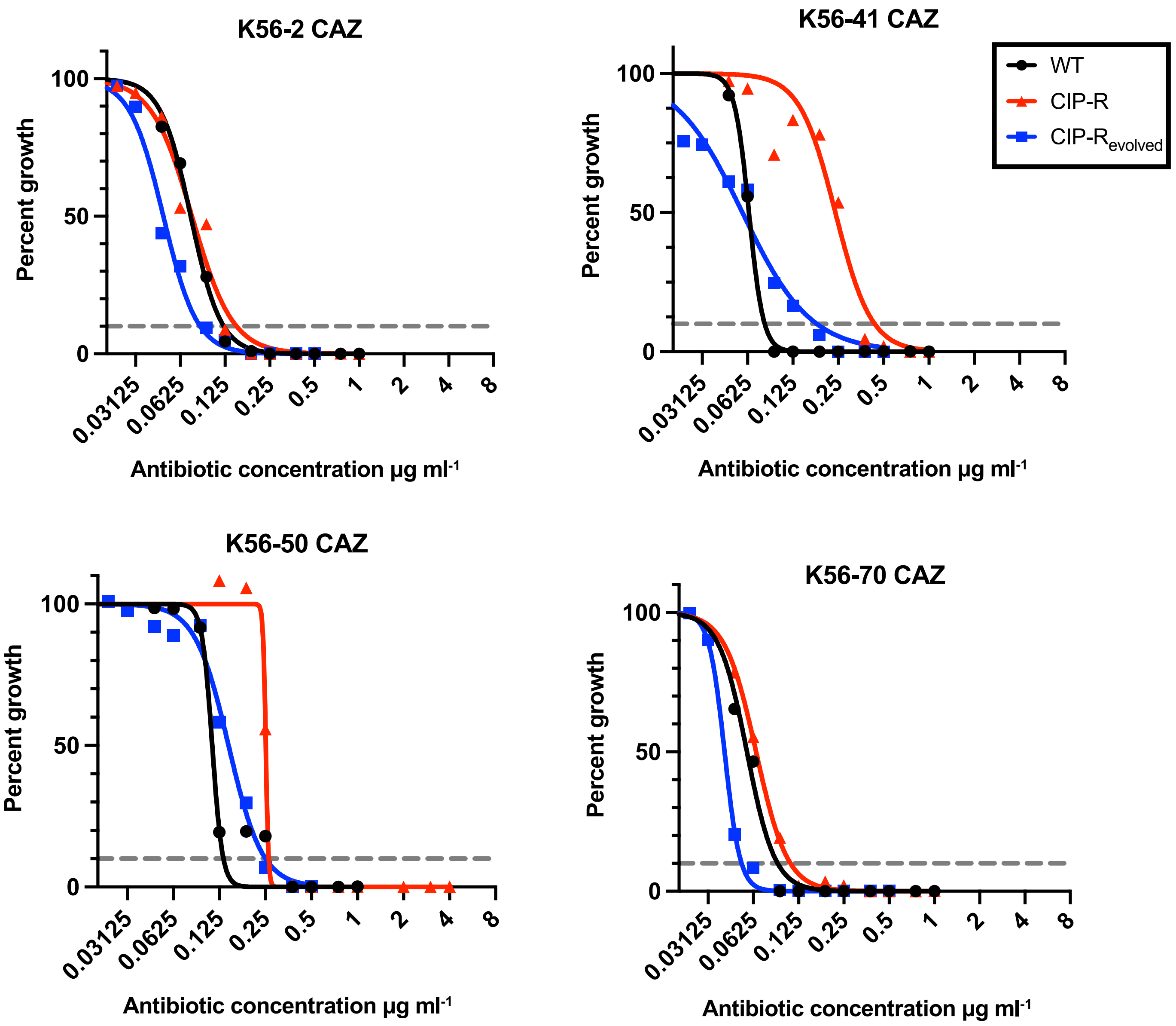


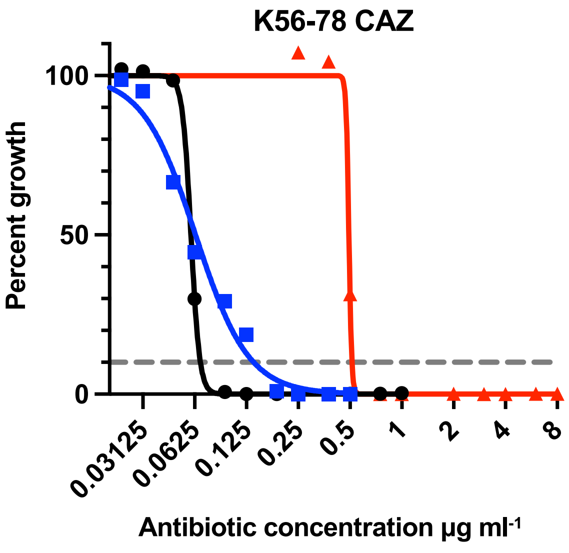


**Supplementary figure 2 – Ceftazidime:**
Dose response curves denoting percent growth at given antibiotic concentrations for Ancestral WT (black circles), Ancestral ciprofloxacin resitant (red triangles) and Evolved ciprofloxacin restant (blue squares). Each strain background are separated into different graphs. Grey dashed line denotes the $IC_{90}$.

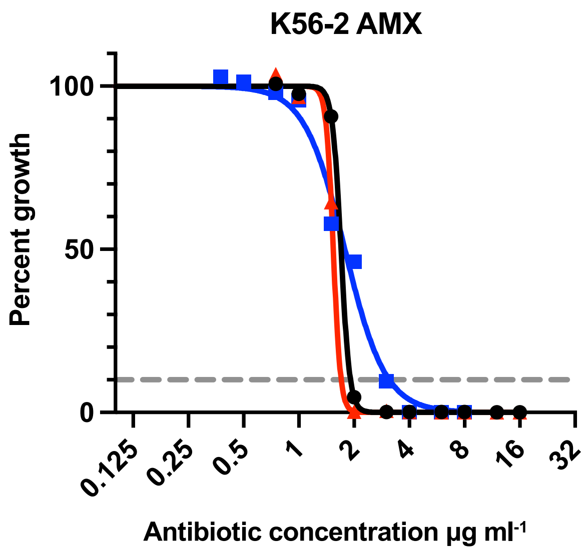


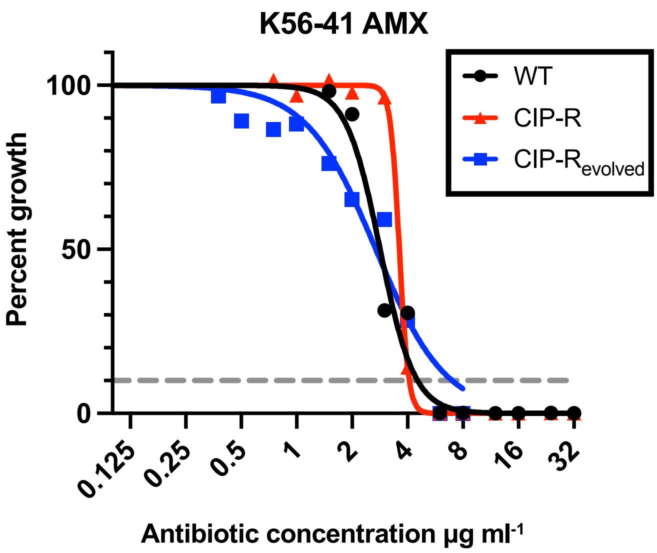


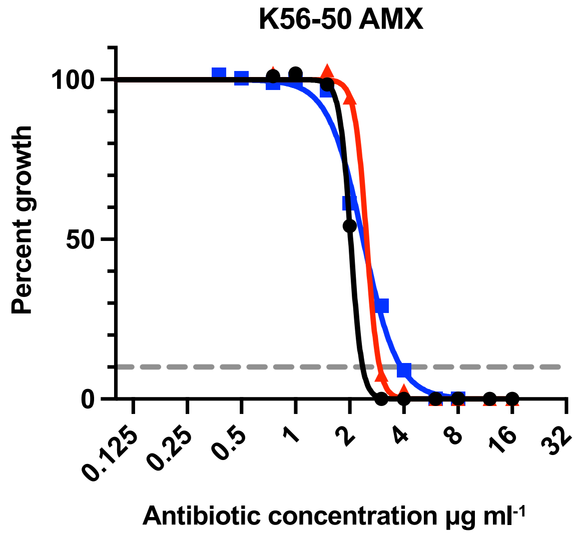


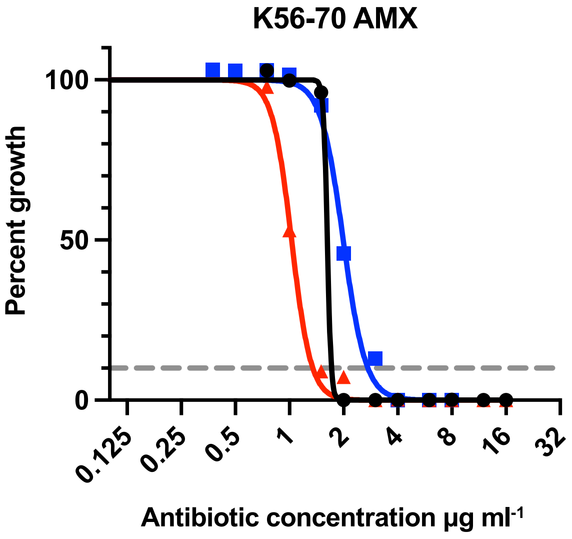


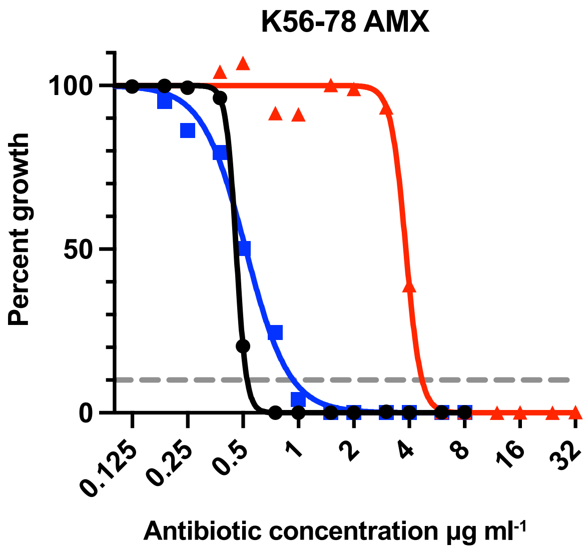


**Supplementary figure 2 – Amoxicillin:**
Dose response curves denoting percent growth at given antibiotic concentrations for Ancestral WT (black circles), Ancestral ciprofloxacin resitant (red triangles) and Evolved ciprofloxacin restant (blue squares). Each strain background are separated into different graphs. Grey dashed line denotes the $IC_{90}$.

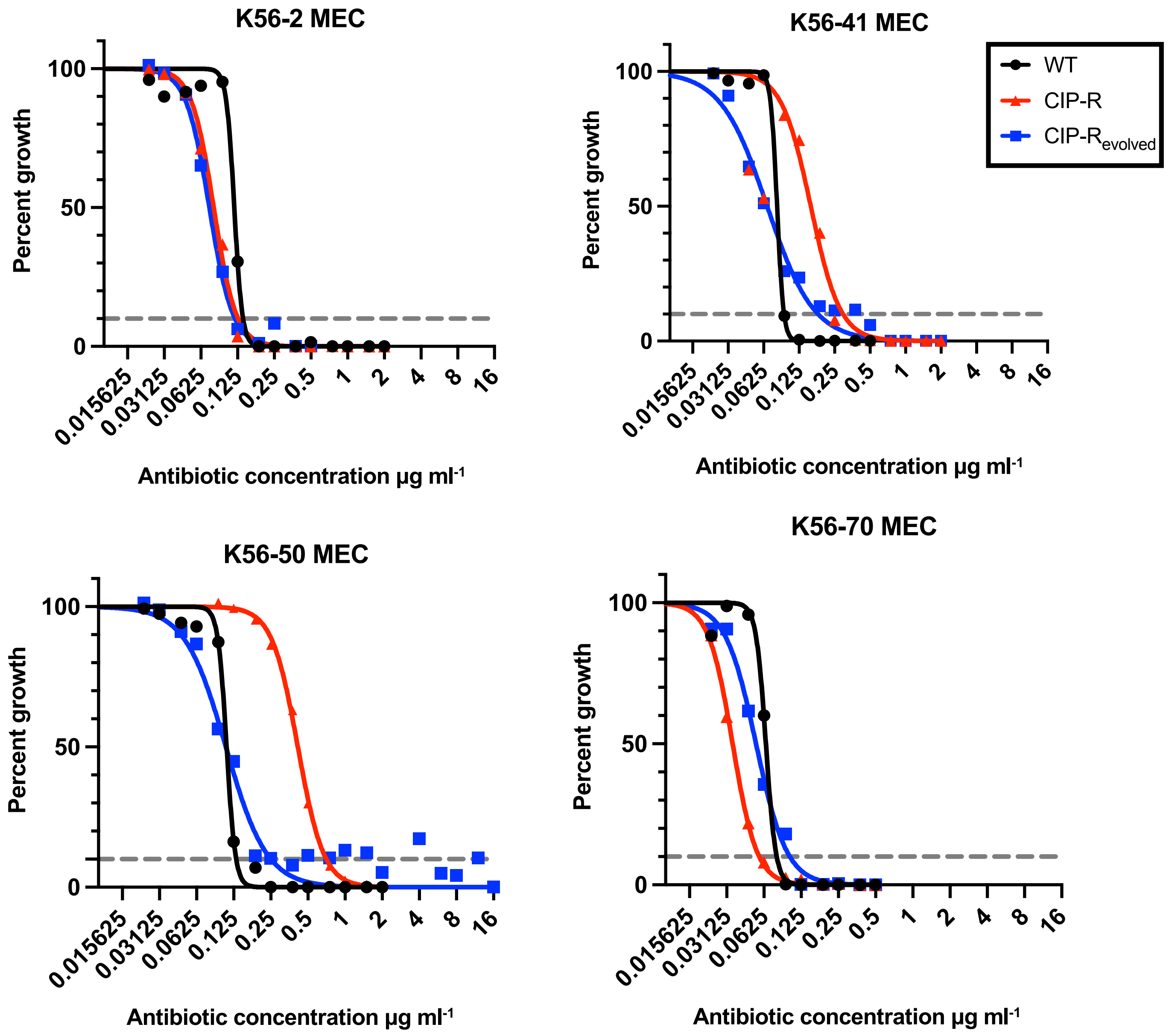


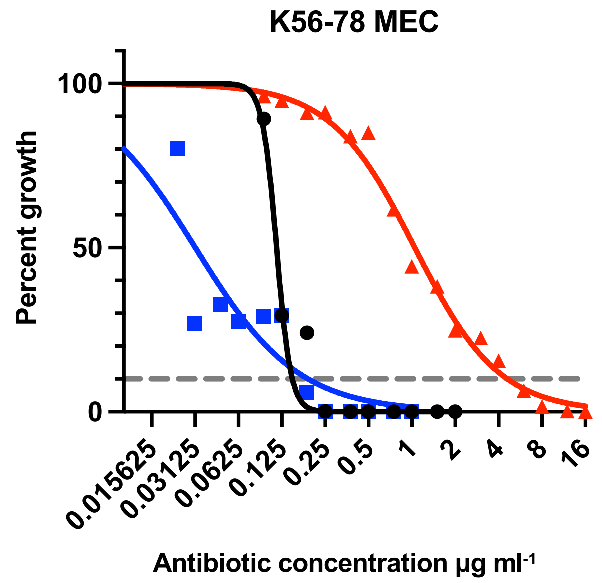


**Supplementary figure 2 – Mecillinam:**
Dose response curves denoting percent growth at given antibiotic concentrations for Ancestral WT (black circles), Ancestral ciprofloxacin resitant (red triangles) and Evolved ciprofloxacin restant (blue squares). Each strain background are separated into different graphs. Grey dashed line denotes the $IC_{90}$.

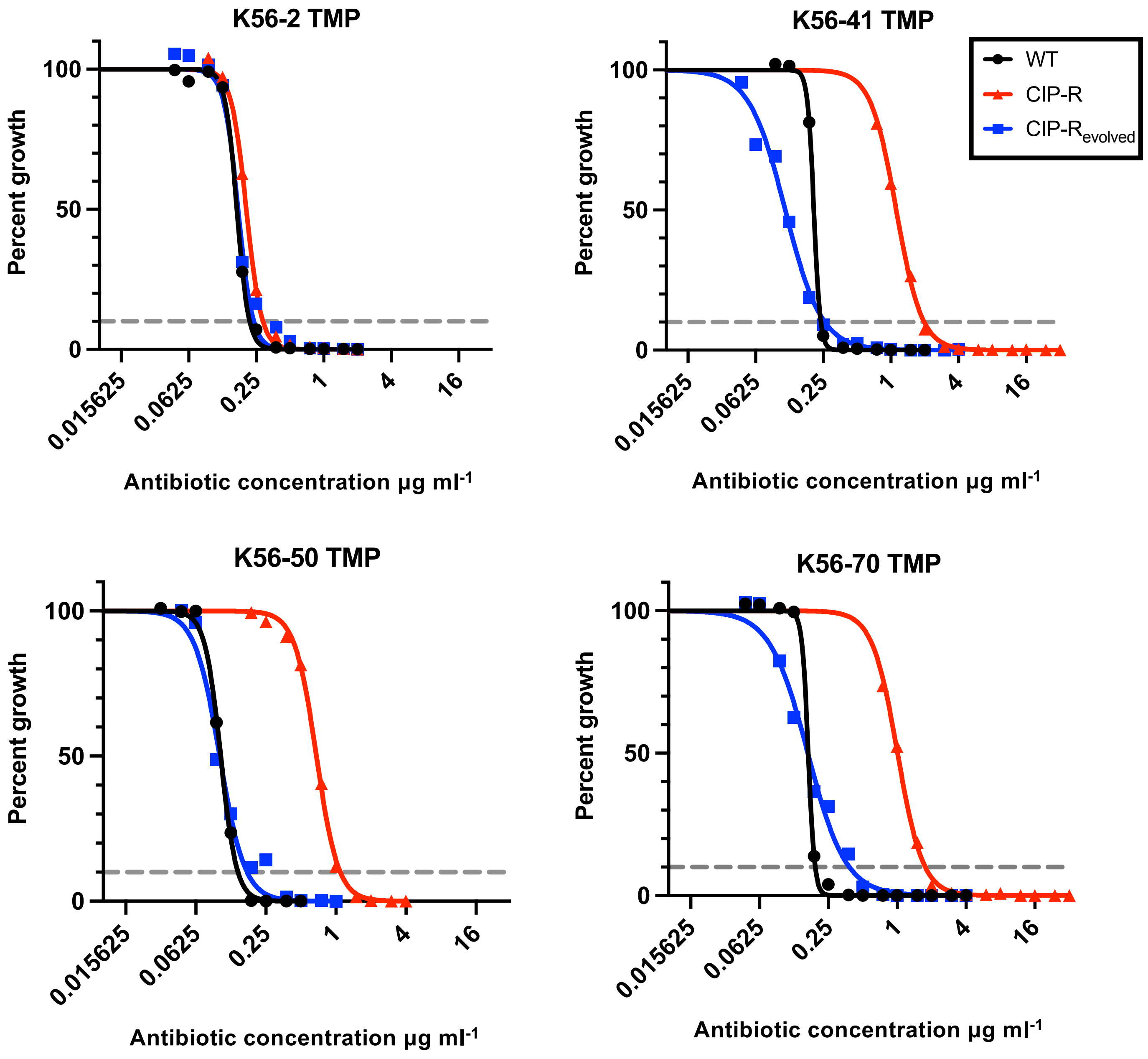


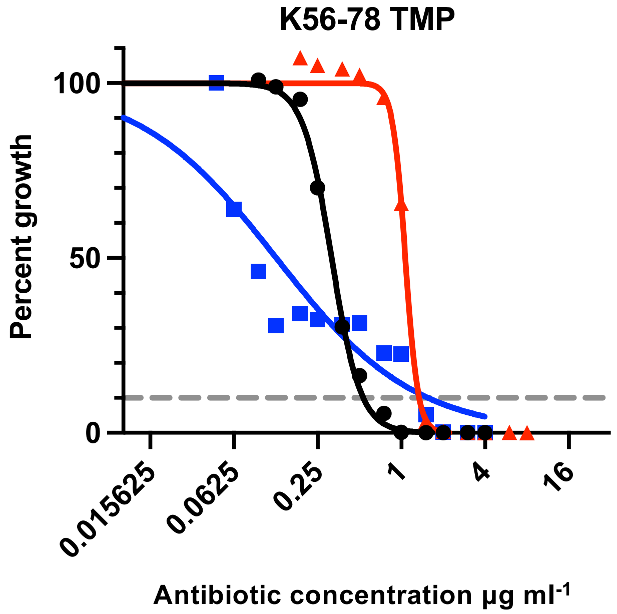


**Supplementary figure 2 – Trimethoprim:**
Dose response curves denoting percent growth at given antibiotic concentrations for Ancestral WT (black circles), Ancestral ciprofloxacin resitant (red triangles) and Evolved ciprofloxacin restant (blue squares). Each strain background are separated into different graphs. Grey dashed line denotes the $IC_{90}$.

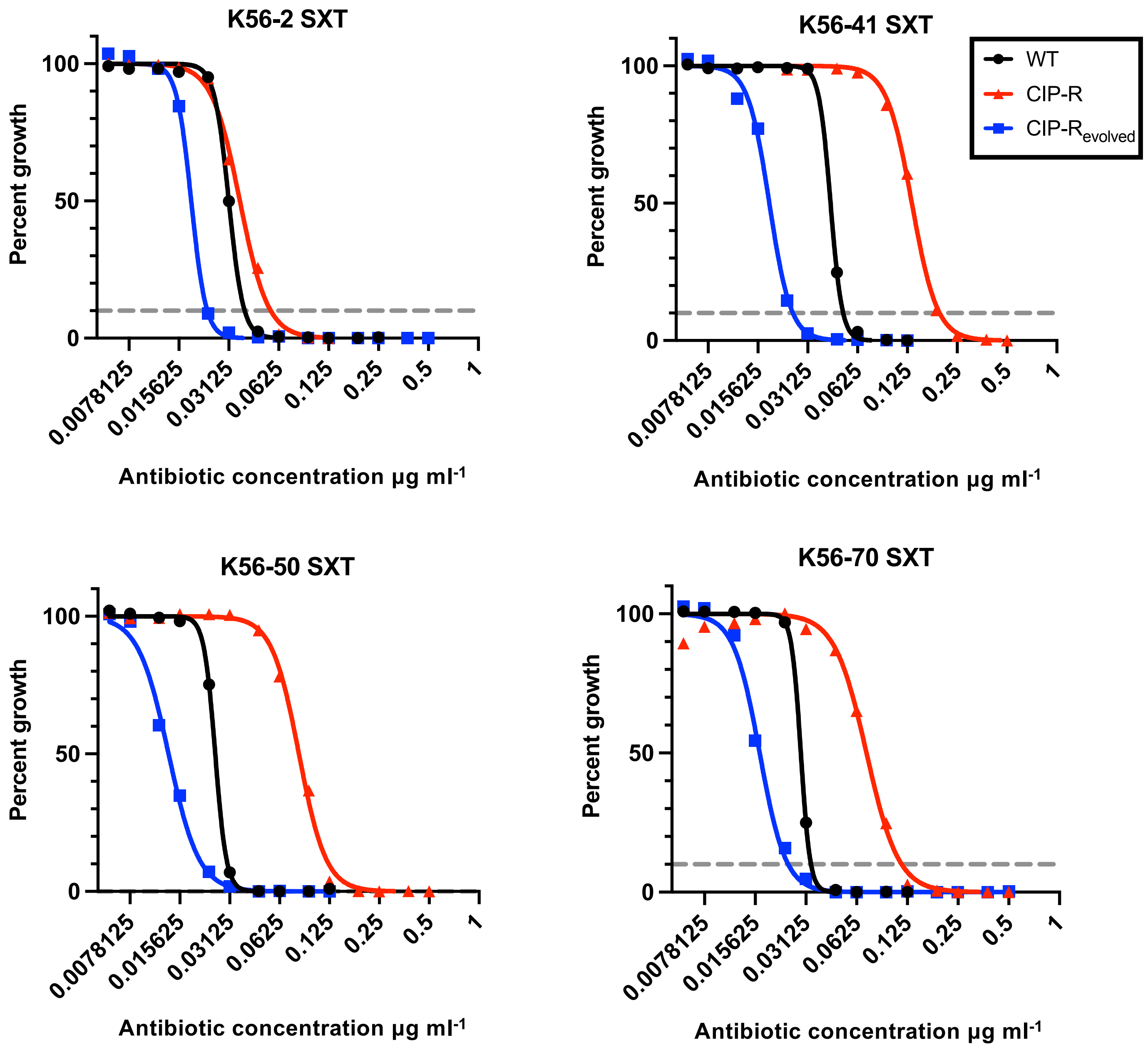


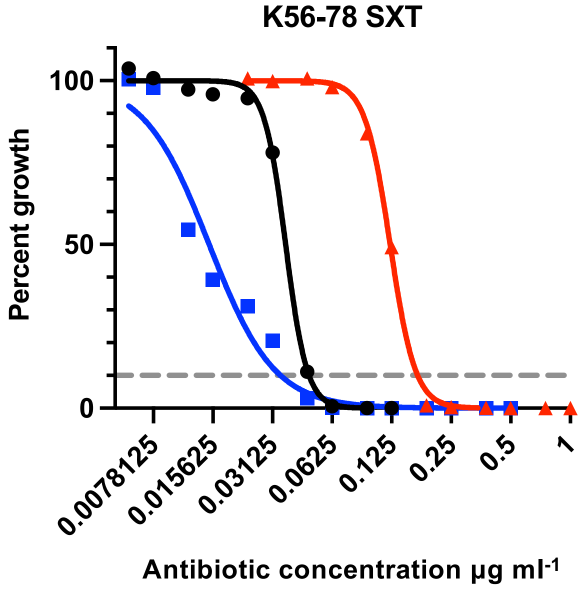


**Supplementary figure 2 – Trimethoprim-Sulfametoxazole:**

Dose response curves denoting percent growth at given antibiotic concentrations for Ancestral WT (black circles), Ancestral ciprofloxacin resitant (red triangles) and Evolved ciprofloxacin restant (blue squares). Each strain background are separated into different graphs. Grey dashed line denotes the $IC_{90}$.

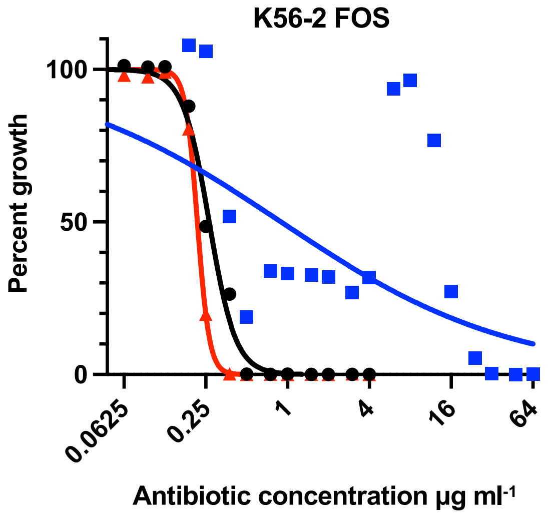


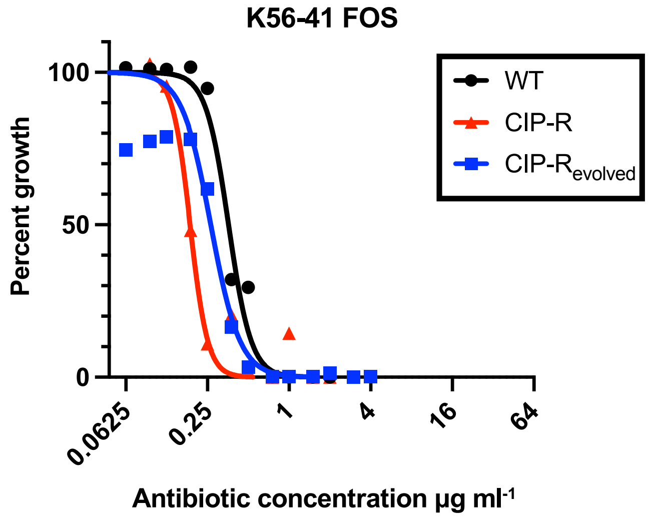


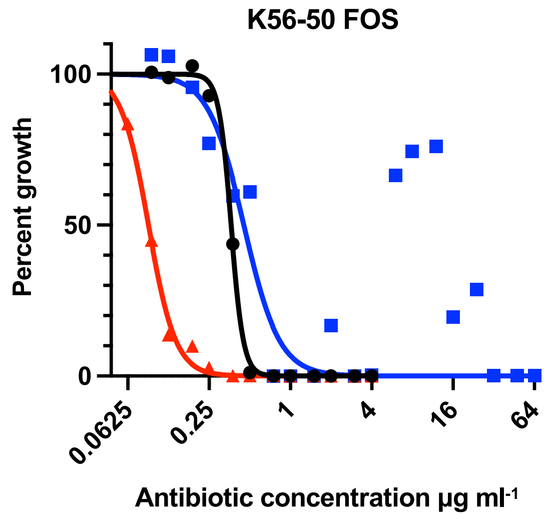


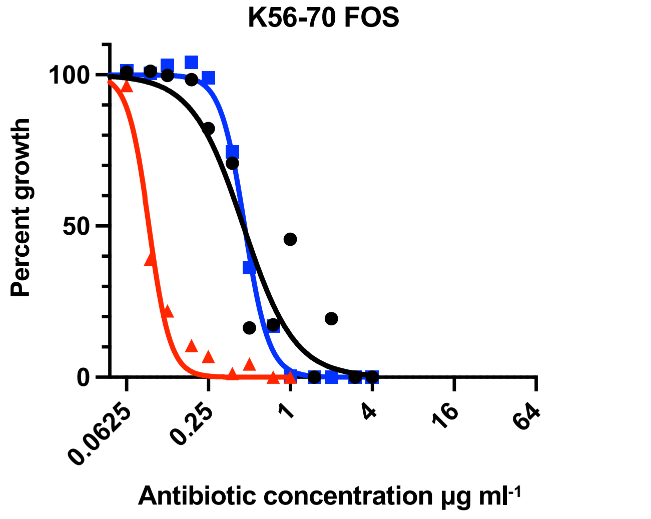


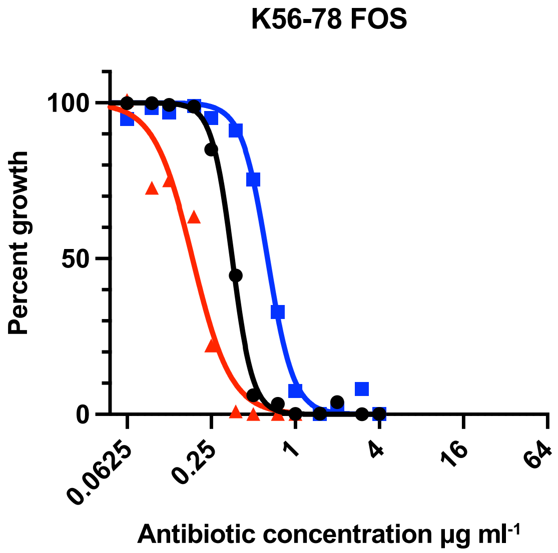


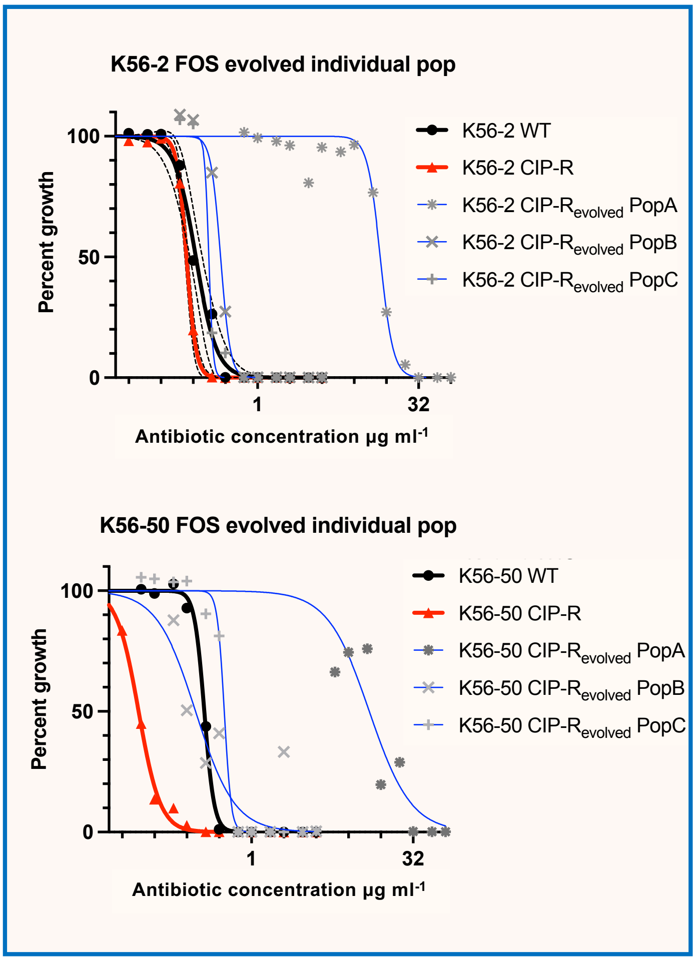


**Supplementary figure 2 – Fosfomycin:**
Dose response curves denoting percent growth at given antibiotic concentrations for Ancestral WT (black circles), Ancestral ciprofloxacin resitant (red triangles) and Evolved ciprofloxacin restant (blue squares). Each strain background are separated into different graphs. Framed in the lower right corner the three individually evolved populations (grey symbols and blue lines) for strain K56-2 and K56-50 is shown separately due to large variation between the populations. Grey dashed line denotes the $IC_{90}$.

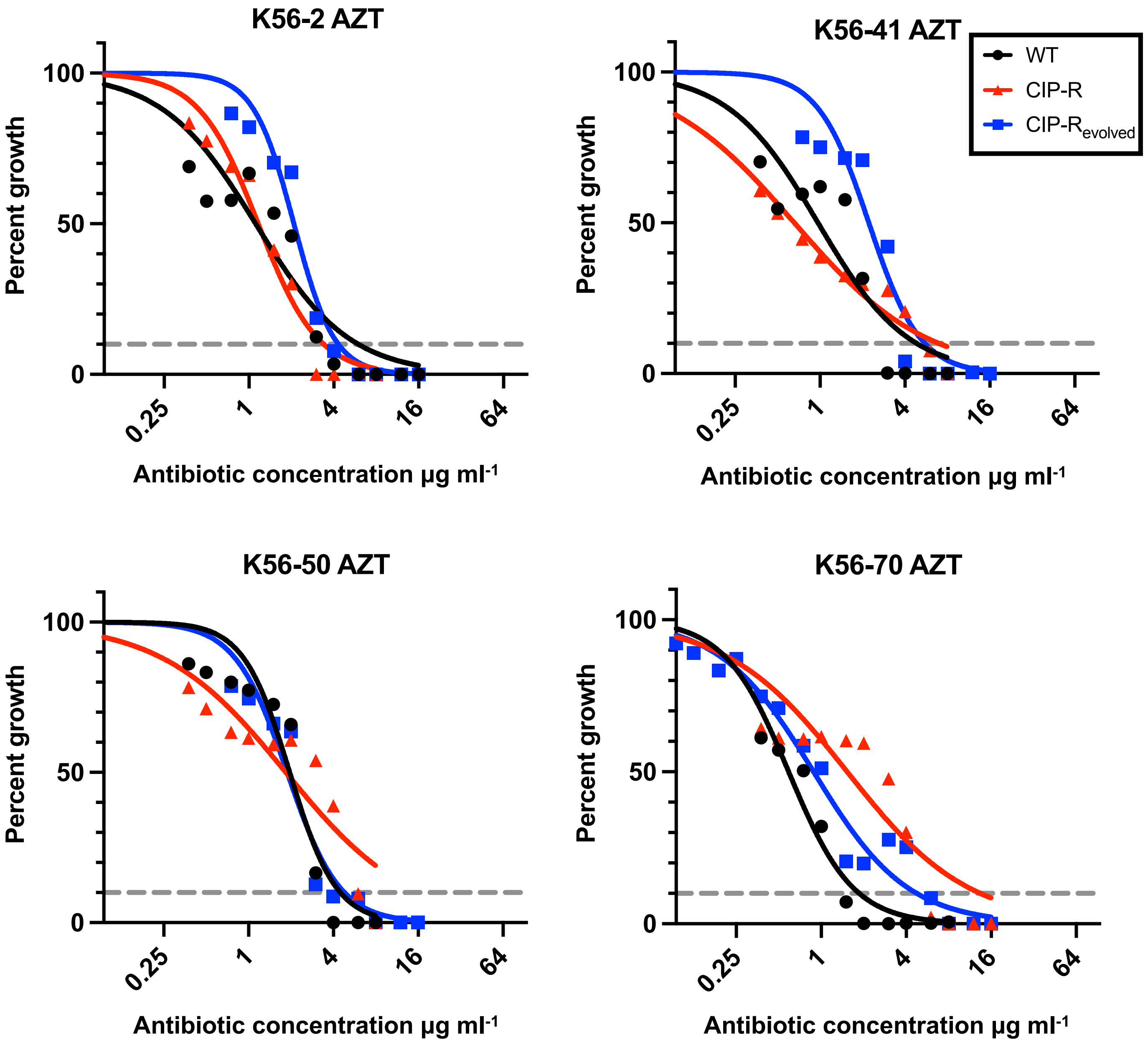


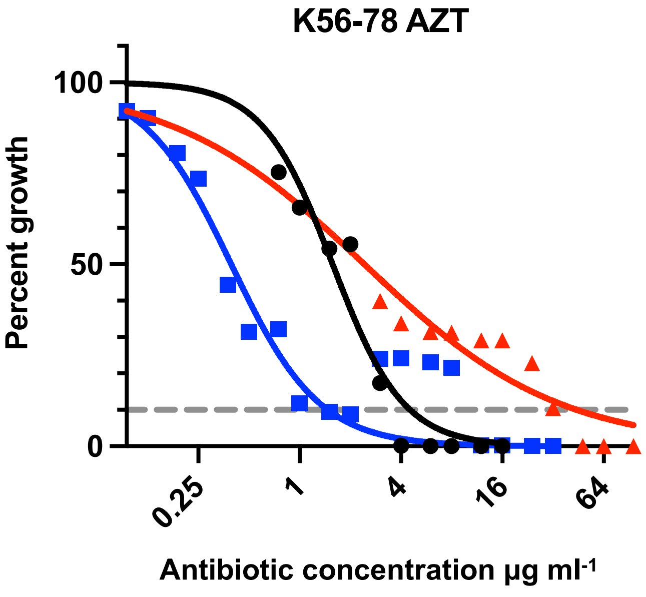


**Supplementary figure 2 – Azithromycin:**
Dose response curves denoting percent growth at given antibiotic concentrations are provided for Ancestral WT (black circles), Ancestral ciprofloxacin resitant (red triangles) and Evolved ciprofloxacin restant (blue squares). The different strain backgrounds are separated into different graphs. Grey dashed line denotes the $IC_{90}$.

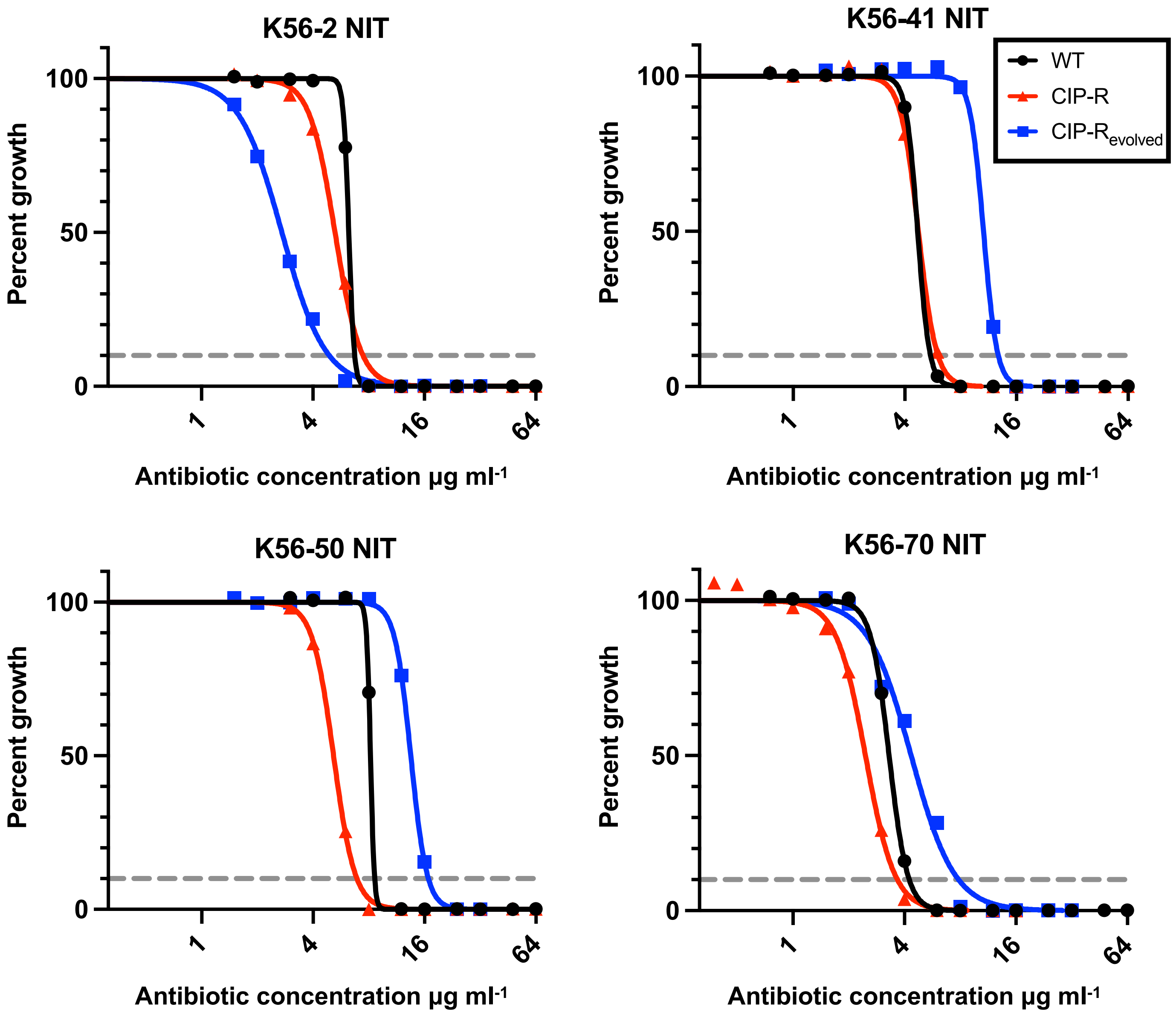


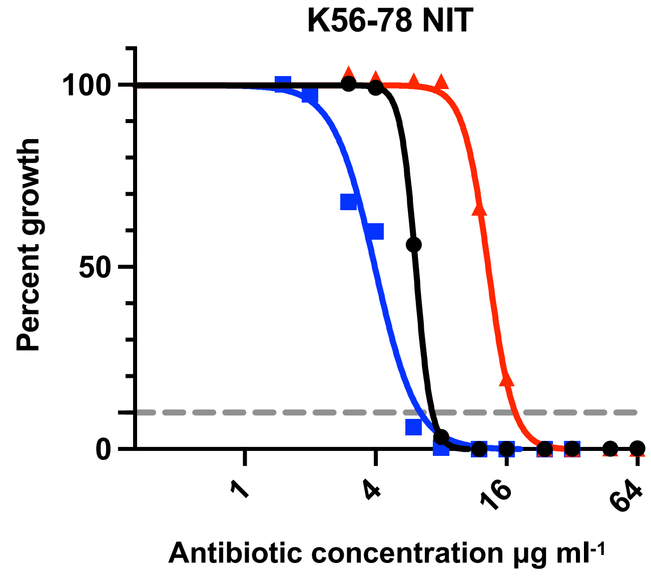


**Supplementary figure 2 – Nitrofurantoin:**
Dose response curves denoting percent growth at given antibiotic concentrations for Ancestral WT (black circles), Ancestral ciprofloxacin resitant (red triangles) and Evolved ciprofloxacin restant (blue squares). Each strain background are separated into different graphs. Grey dashed line denotes the $IC_{90}$.

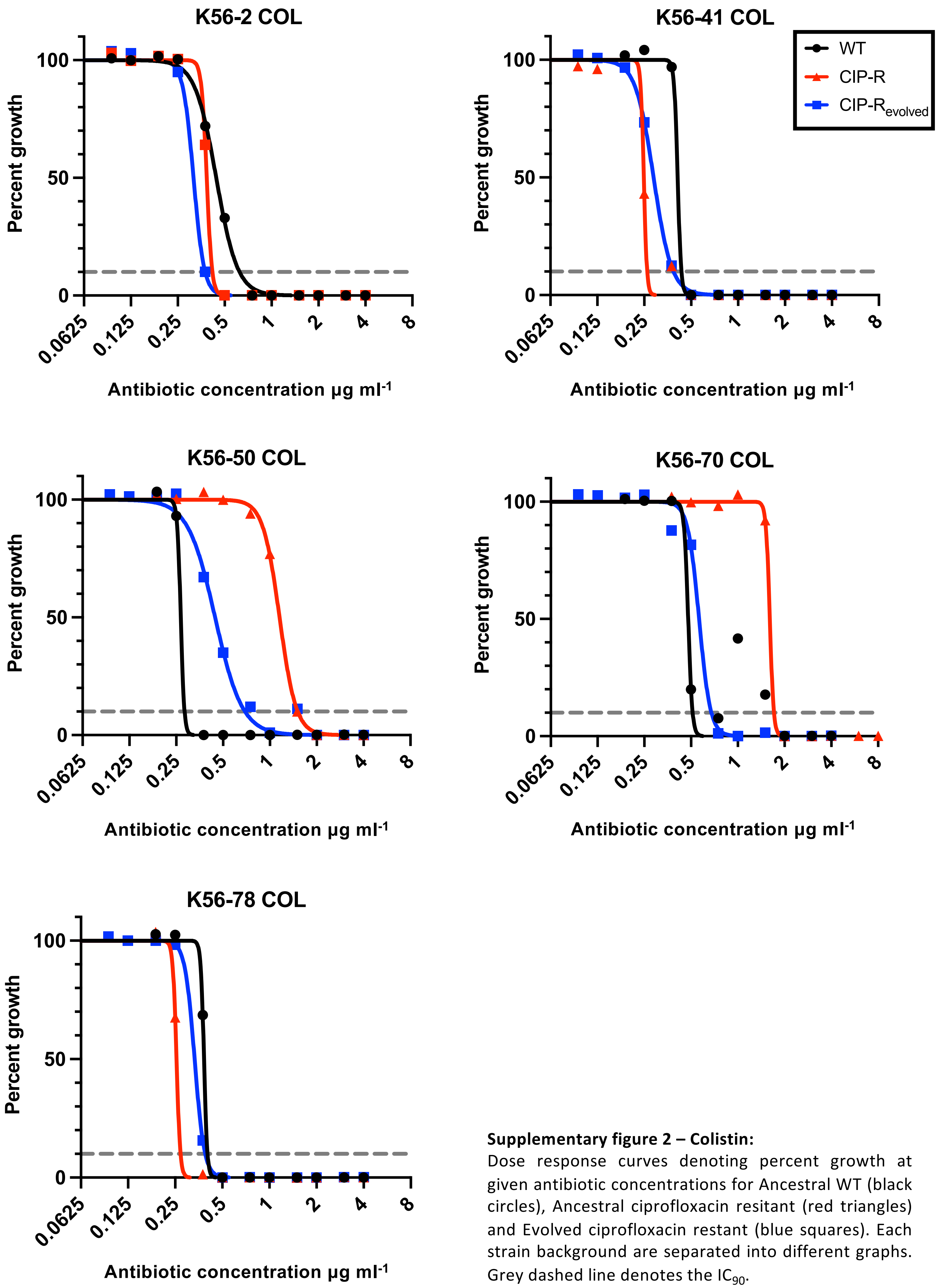


**Supplementary figure 2 – Colistin:**
Dose response curves denoting percent growth at given antibiotic concentrations for Ancestral WT (black circles), Ancestral ciprofloxacin resitant (red triangles) and Evolved ciprofloxacin restant (blue squares). Each strain background are separated into different graphs. Grey dashed line denotes the $IC_{90}$.

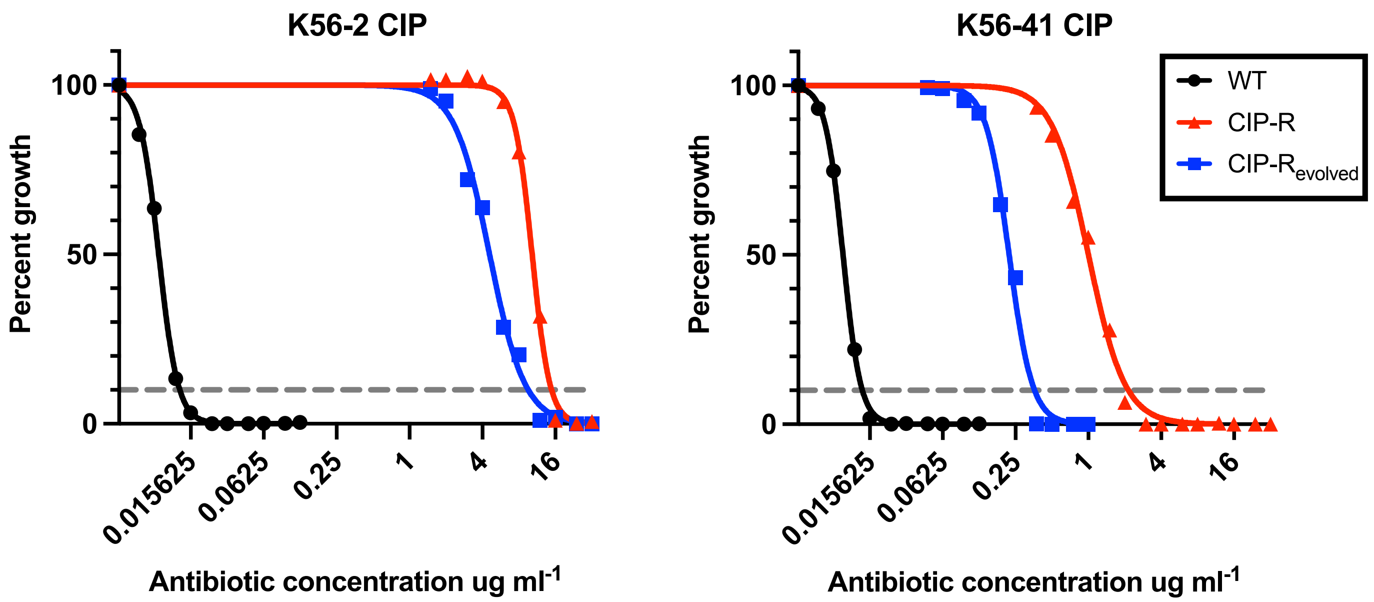


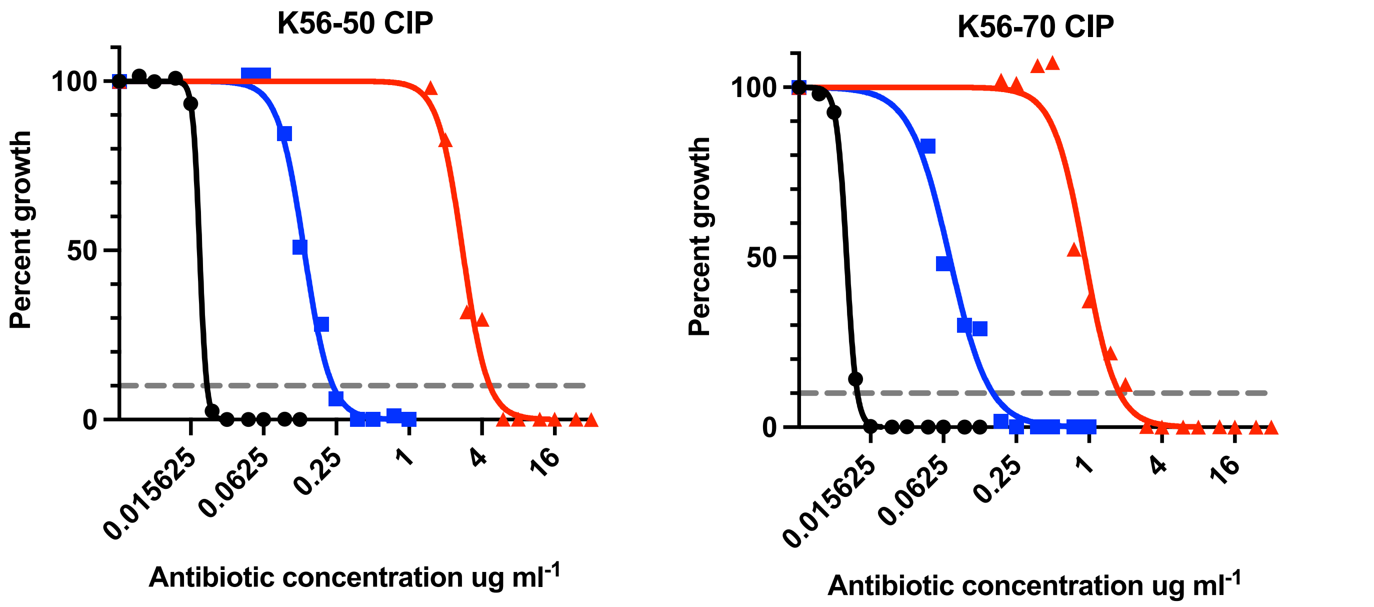


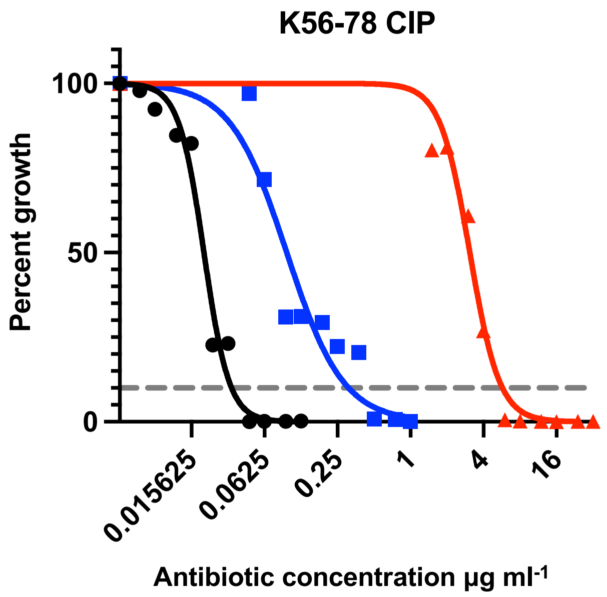


**Supplementary figure 2 – Ciprofloxacin:**
Dose response curves denoting percent growth at given antibiotic concentrations for Ancestral WT (black circles), Ancestral ciprofloxacin resitant (red triangles) and Evolved ciprofloxacin restant (blue squares). Each strain background are separated into different graphs. Grey dashed line denotes the $IC_{90}$.

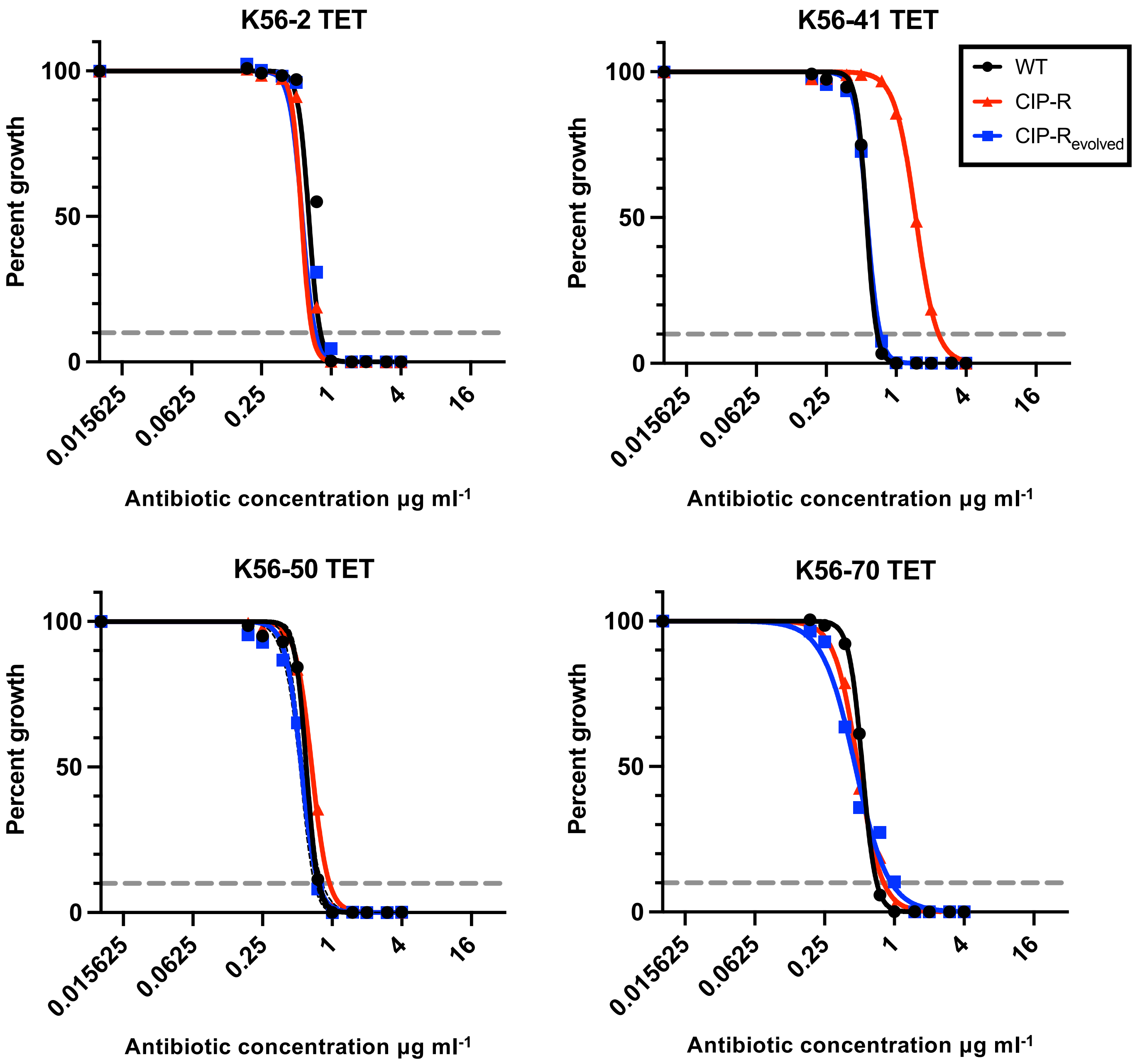


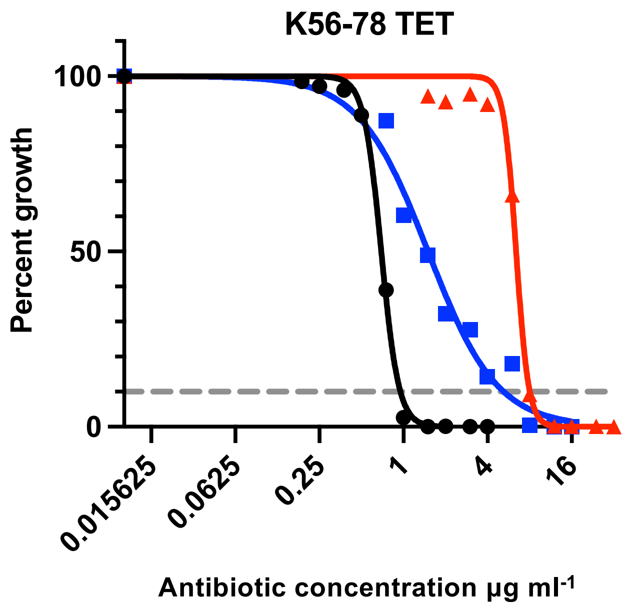


**Supplementary figure 2 – Tetracycline:**
Dose response curves denoting percent growth at given antibiotic concentrations for Ancestral WT (black circles), Ancestral ciprofloxacin resitant (red triangles) and Evolved ciprofloxacin restant (blue squares). Each strain background are separated into different graphs. Grey dashed line denotes the $IC_{90}$.

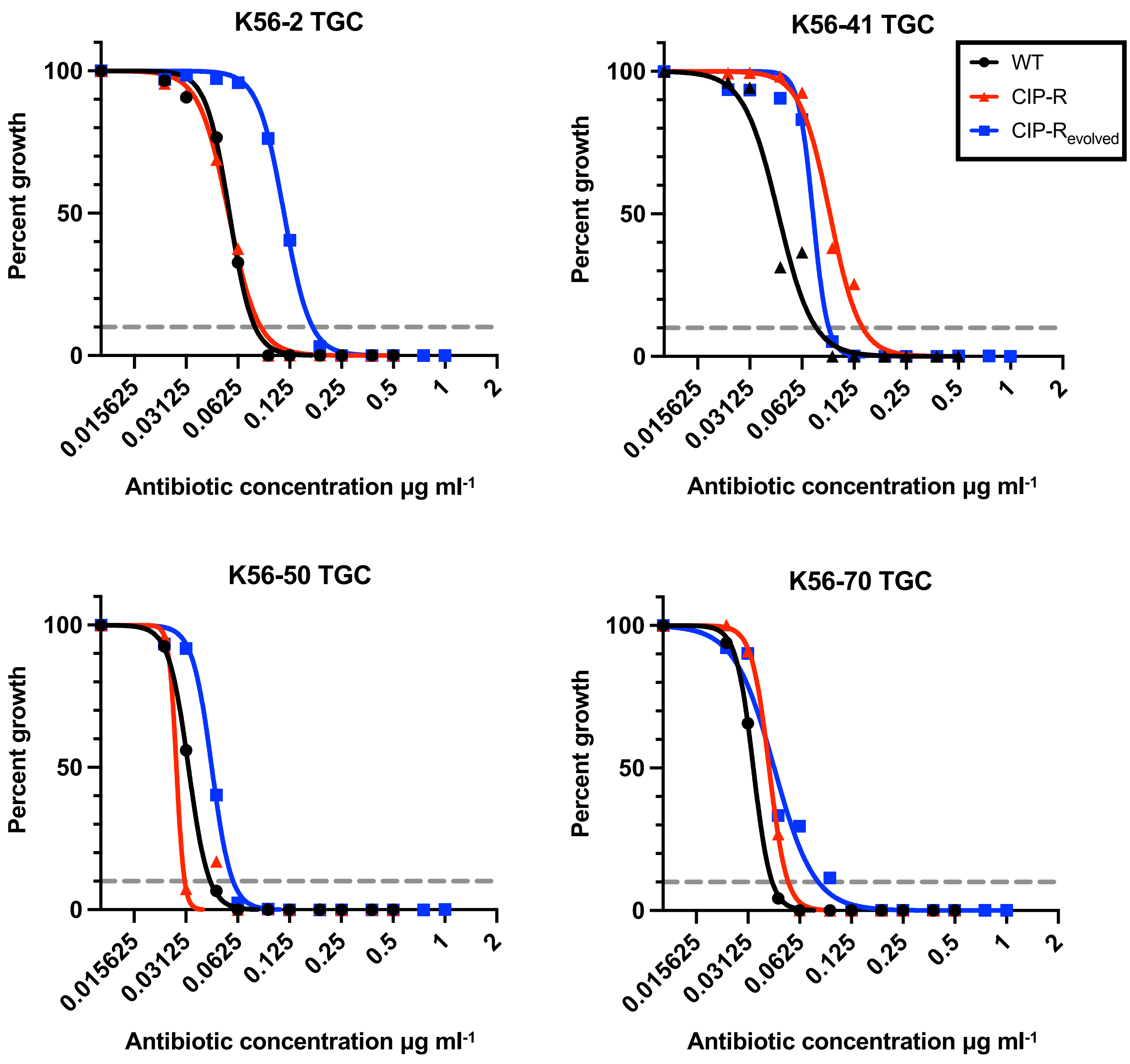


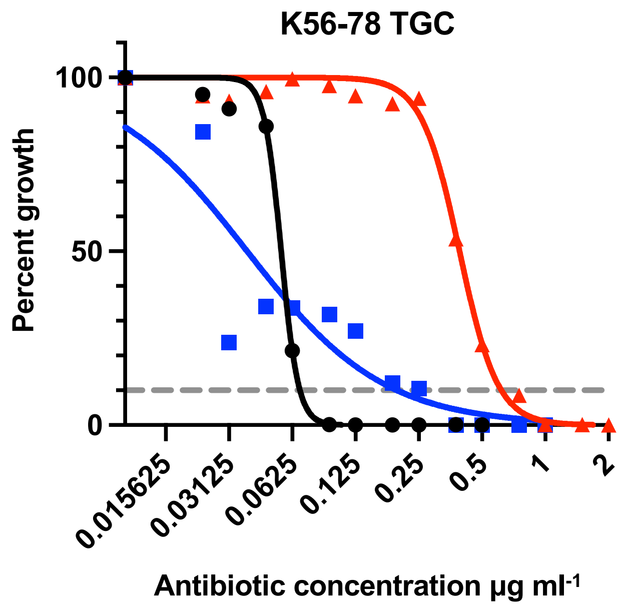


**Supplementary figure 2 – Tigecycline:**
Dose response curves denoting percent growth at given antibiotic concentrations for Ancestral WT (black circles), Ancestral ciprofloxacin resitant (red triangles) and Evolved ciprofloxacin restant (blue squares). Each strain background are separated into different graphs. Grey dashed line denotes the $IC_{90}$.
